# Unisexual reproduction promotes competition for mating partners in the global human fungal pathogen *Cryptococcus deneoformans*

**DOI:** 10.1101/523894

**Authors:** Ci Fu, Torin P. Thielhelm, Joseph Heitman

## Abstract

Courtship is pivotal for successful mating. However, courtship is challenging for the *Cryptococcus neoformans* species complex, comprised of opportunistic fungal pathogens, as the majority of isolates are α mating type. In the absence of mating partners of the opposite mating type, *C. deneoformans* can undergo unisexual reproduction, during which a yeast-to-hyphal morphological transition occurs. Hyphal growth during unisexual reproduction is a quantitative trait, which reflects a strain’s ability to undergo unisexual reproduction. In this study, we determined whether unisexual reproduction confers an ecological benefit by promoting foraging for mating partners. Through competitive mating assays using strains with different abilities to produce hyphae, we showed that unisexual reproduction potential did not enhance competition for mating partners of the same mating type, but when cells of the opposite mating type were present, cells with enhanced hyphal growth were more competitive for mating partners of either the same or opposite mating type. Enhanced mating competition was also observed in a strain with increased hyphal production that lacks the mating repressor gene *GPA3*, which contributes to the pheromone response. Hyphal growth in unisexual strains also enables contact between adjacent colonies and enhances mating efficiency during mating confrontation assays. The pheromone response pathway activation positively correlated with unisexual reproduction hyphal growth during bisexual mating and exogenous pheromone promoted bisexual cell fusion. Despite the benefit in competing for mating partners, unisexual reproduction conferred a fitness cost. Taken together, these findings suggest *C. deneoformans* employs hyphal growth to facilitate contact between colonies at long distances and utilizes pheromone sensing to enhance mating competition.

**Author Summary:** Sexual reproduction plays a pivotal role in shaping fungal population structure and diversity in nature. The global human fungal pathogen *Cryptococcus neoformans* species complex evolved distinct sexual cycles: bisexual reproduction between mating partners of the opposite mating types, and unisexual reproduction with only one mating type. During both sexual cycles, cells undergo a yeast-to-hyphal morphological transition and nuclei diploidize through either cell-cell fusion followed by nuclear fusion during bisexual reproduction or endoreplication during unisexual reproduction. Despite the complex sexual life cycle, the majority of Cryptococcal isolates are α mating type. Albeit the scarcity of *MAT***a** cells in the environment, meiotic recombination is prevalent. To decipher this conundrum, we ask whether there is an underlying mechanism in which *Cryptococcus* species increase their mating opportunities. In this study, we showed that the undirected hyphal growth during unisexual reproduction enables *MAT*α cells to forage for mating partners over a larger surface area, and when *MAT*α hyphae come into close proximity of rare *MAT***a** cells, pheromone response pathway activation in both *MAT*α and *MAT***a** cells can further enhance mating. This mating enhancement could promote outcrossing and facilitate genome reshuffling via meiotic recombination.

## Introduction

Successful courtship is key to the evolutionary success of sexual organisms, and many species have evolved distinct strategies to locate and choose a mating partner. For example, primates and humans utilize aggression to secure a mating partner [1]; male hummingbirds apply acoustic control using tail feathers during high-speed dives to court females [2]; male *Drosophila* vibrate their wings to generate different songs to trigger mating responses in females [3]; male tree-hole frogs also adopt acoustic strategies taking advantage of tree trunk cavities to attract females [4]; and female pipefish display a temporal striped pattern ornament to woo male partners [5]. These examples demonstrate that complex eukaryotic organisms can employ visual, vocal, or mechanical tactics to secure a mate and transmit their genetic traits to the next generation.

In eukaryotic fungal systems, mating often involves a morphological transition. *Saccharomyces cerevisiae* yeast cells undergo polarized growth and form shmoo projections in preparation for cell fusion during mating [6]. In filamentous fungi, including both ascomycetes and basidiomycetes, sexual reproduction involves the formation of a fruiting body (perithecium or basidium, respectively) [7]. *Candida albicans*, an ascomycete, undergoes a white-opaque switch to initiate mating [8]. Despite their divergent sexual strategies, these morphological transitions are all controlled by the pheromone response pathway [9]. During yeast mating, physical agglutination of yeast cells does not promote courtship, but rather a gradient of pheromone signals is crucial for successful cell-cell fusion during early mating [10, 11]. Similarly, in *Schizosaccharomyces pombe*, local pheromone signals and a spatially focal pheromone response dictate cell-cell pairing and fusion position during early mating processes [12, 13]. In *C. albicans*, overexpression of the pheromone response MAP kinase pathway components can enhance mating efficiency [14]. These studies establish that the pheromone response pathway plays a critical role in promoting fungal mating efficiency.

The opportunistic human fungal pathogen *Cryptococcus deneoformans* undergoes a yeast-to-hyphal morphological transition upon mating induction [15]. This species has two modes of sexual reproduction: bisexual reproduction between cells of opposite mating types and unisexual reproduction involving cells of only one mating type [15-17]. Cell fusion between *MAT***a** and *MAT*α cells during bisexual reproduction, and between two *MAT*α cells during unisexual reproduction, triggers hyphal development [18]. This morphological transition is orchestrated by the pheromone response pathway [18, 19]. However, recent studies have shown that hyphal growth during unisexual reproduction can also occur independent of cell fusion and the pheromone response pathway [20-23], and that pheromone-independent hyphal development is dependent upon the calcineurin pathway [20, 24].

Because the majority of identified natural and clinical *C. neoformans* isolates are of the α mating type, unisexual reproduction likely has significant ecological impacts on the *Cryptococcus* species complex population structure and diversity [25-27]. The limited abundance of *MAT***a** cells in natural environments restricts outcrossing and in the absence of **a**-α mating, unisexual reproduction has been shown to reverse Muller’s rachet and offset the low abundance of *MAT***a** cells to avoid an evolutionary dead end [28]. Unisexual reproduction can also generate genotypic and phenotypic diversity *de novo* [29]. Interestingly, population genetics studies have revealed that genome recombination occurs frequently among environmental isolates [30-32], even those that are exclusively α mating type, providing evidence that unisexual reproduction involving fusion of *MAT*α cells of distinct genotypes allows meiotic recombination in nature. Despite these evolutionary benefits, cell fusion-independent solo-unisexual reproduction also occurs and because this pathway involves genetically identical genomes, it does not contribute to genome reshuffling or recombination. Similar to pseudohyphal differentiation in *S. cerevisiae, C. deneoformans* hyphal growth during unisexual reproduction has an ecological benefit in promoting foraging for nutrients and habitat exploration in the surrounding environments [33, 34]. In this study, we address whether the ability to undergo unisexual reproduction has an additional ecological benefit in promoting foraging for mating partners to facilitate outcrossing and enable recombination in nature.

## Results and Discussion

### Strains with enhanced unisexual reproduction potential are more competitive for mating partners of the opposite mating type

During *C. deneoformans* solo-unisexual reproduction, cells undergo the yeast-to-hyphal morphological transition independent of cell fusion and nuclei diploidized through endoreplication [16, 23]. The hyphal growth is a quantitative trait associated with unisexual reproduction that can be used to determine a strain’s ability to undergo unisexual reproduction [35]. Although solo-unisexual reproduction occurs independently of cell-cell fusion, cells can fuse with partners of both the same or opposite mating type at varying frequencies [16, 23]. To test whether the ability to undergo unisexual reproduction impacts competition for mating partners during outcrossing, we performed mating competition experiments employing three *MAT*α and three *MAT***a** *C. deneoformans* strains with different degrees of unisexual reproduction potential based on their abilities to produce hyphae (Figure 1A) [35]. Among these strains, several were F2 progeny derived from crosses between the environmental *MAT***a** isolate NIH433 and the clinical *MAT*α isolate NIH12 including a high hyphal (HH) strain XL190α, an intermediate hyphal (MH) strain XL280α, a low hyphal strain XL187**a**, and a no hyphal (NH) strain JEC20**a** [15, 16, 36-38]. LH strain JEC21α and MH strain XL280**a** are congenic strains of JEC20**a** and XL280α, respectively, derived through 10 rounds of backcrossing (Figure S1) [36, 38, 39]. For each mating competition experiment, cells of three strains with different hyphal growth carrying dominant, selectable drug resistance markers were mixed, spot-inoculated, and incubated on V8 agar media for 4 days (Figure 1B). Cells were recovered on YPD medium to obtain colony forming units (CFU), and on YPD medium supplemented with different two-drug combinations to determine the cell fusion frequencies. Cell fusion frequencies were compared between different pairs of strains within the same competition mating mixture to determine whether the ability to undergo unisexual reproduction confers benefits in competition for mating partners to facilitate outcrossing (Figure 1B).

**Figure 1.**
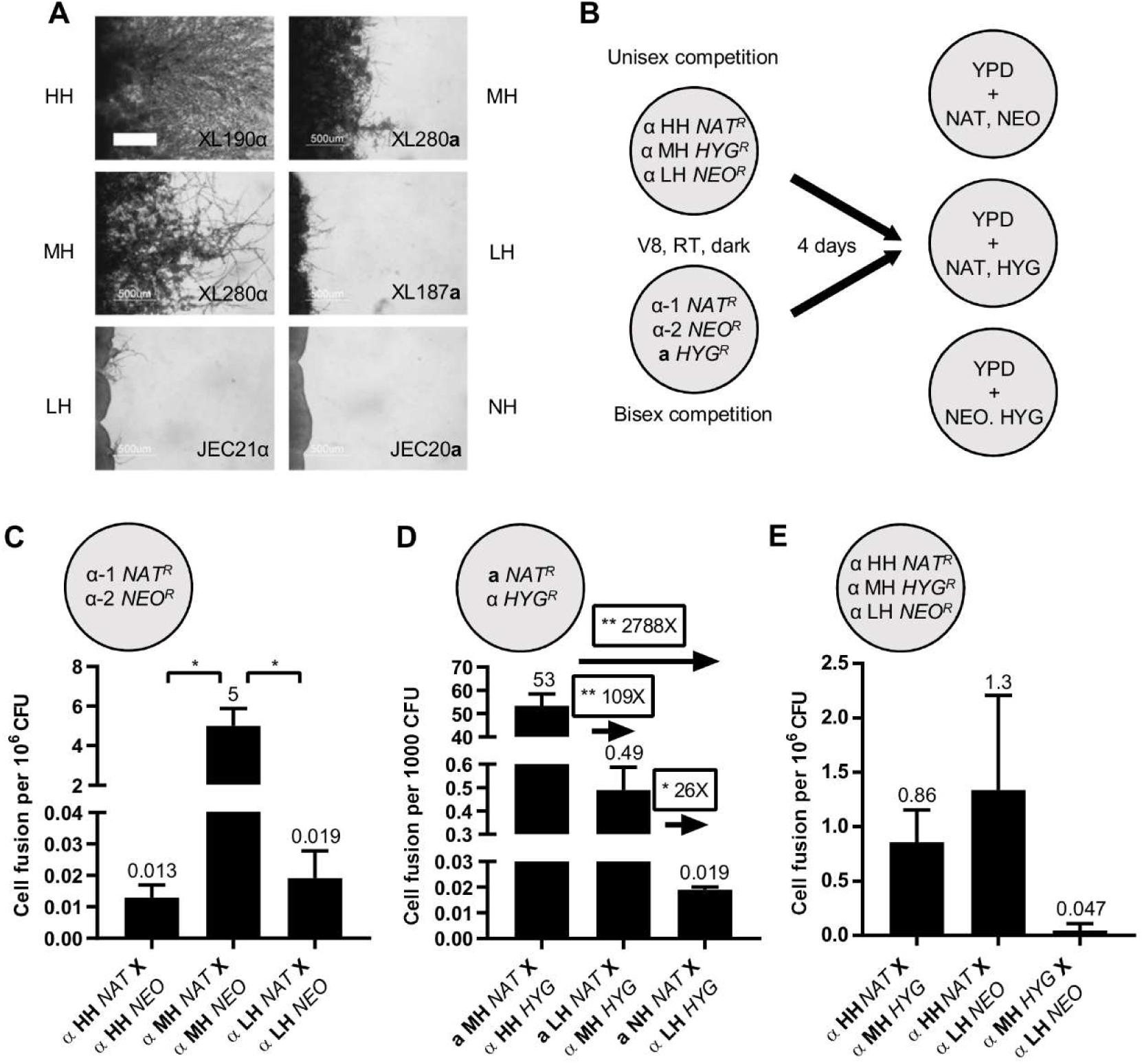
Unisexual reproduction potential does not enhance competition for mating partners of the same mating type. (A) Hyphal growth on V8 agar medium for three weeks of high (HH), intermediate (MH), low (LH), and no (NH) hyphal *MAT*α and *MAT***a** strains. The scale bar represents 500 *µ*m. (B) Experimental design of the unisexual and bisexual mating competition. (C-E) Average cell fusion frequencies. Error bars indicate standard deviation. (C) Unisexual α-α cell fusion frequencies of high, intermediate, and low hyphal strains. (D) Bisexual **a**-α cell fusion frequencies of intermediate-high, low-intermediate, and no-low hyphal strains. (* indicates 0.01<*p*≤0.05 and ** indicates 0.001<*p*≤0.01 for each pairwise comparison) (E) Cell fusion frequencies among three *MAT*α strains of different hyphal growth phenotypes during unisexual mating competition are shown.

Prior to the mating competition experiments, cell fusion frequencies were compared between different hyphal strains. During α-α cell fusion, the *MAT*α MH strain displayed a significantly higher cell fusion frequency (5 cell fusion events per million CFU) compared to the HH and LH strains (0.013 and 0.019 cell fusion events per million CFU, respectively), in which cell fusion rarely occurred (Figure 1C). This suggests that the ability to undergo more robust hyphal growth is not strictly correlated with α-α cell fusion efficiency. In contrast, during **a**-α cell fusion, hyphal growth positively correlated with **a**-α cell fusion efficiency. MH-HH strains had a cell fusion frequency (53 cell fusion events per thousand CFU) about 109 times higher than LH-MH strains, which in turn had a cell fusion frequency (0.49 cell fusion events per thousand CFU) about 26 times higher than NH-LH strains (0.019 cell fusion events per thousand CFU) (Figure 1D). In all of the strains tested, **a**-α cell fusion occurred at a much higher level compared to α-α cell fusion, similar to previous findings [16, 23].

Cell fusion has been previously shown to be dispensable for solo-unisexual reproduction [19, 23], which can account for the observed poor correlation between hyphal growth and α-α cell fusion frequency. Thus, we hypothesize that increased hyphal growth may not provide an advantage in competing for mating partners of the same mating type. Indeed, when we performed the unisexual mating competition assay mixing the HH, MH, and LH cells, we observed that HH and LH cells yielded the most fusion products with a cell fusion frequency of 1.3 cell fusion events per million CFU that is not significantly different from cell fusion frequencies involved MH cells (Figure 1E) that exhibited the highest cell fusion frequency (Figure 1C). These findings indicate that neither α-α cell fusion frequency nor hyphal growth can be used to predict mating partner preference during unisexual reproduction, which supports the hypothesis that the ability to undergo unisexual reproduction does not promote competition for mating partners of the same mating type.

To test whether the propensity for unisexual reproduction plays a role in competing for mating partners of the opposite mating type, mating competition assays were conducted for a given *MAT***a** isolate between two *MAT*α strains of different hyphal growth phenotypes (Figure 2A). Interestingly, cells capable of producing more hyphae always had a significantly higher cell fusion frequency with *MAT***a** cells compared to cells with lower hyphal growth potential (Figure 2A, Table S1). For example, in the presence of MH *MAT***a** cells, HH *MAT*α cells fused with *MAT***a** cells 24 times more efficiently than LH *MAT*α cells and 8.1 times more efficiently than MH *MAT*α cells, and MH *MAT*α cells fused with *MAT***a** cells 5.8 times more efficiently than LH *MAT*α cells (Table S1). These results suggest that increased hyphal growth correlates with competition for mating partners of the opposite mating type during bisexual reproduction. It was also noted that the mating competition advantage decreased for each competition pair (24, 14.5, and 8.9 fold differences for HH vs LH, 8.1, 6.5, and 5.3 fold differences for HH vs MH, and 5.8, 4.6, and 1.7 fold differences for MH vs LH) with the decreasing hyphal phenotype of the *MAT***a** cells (Figure 2A and Table S1), suggesting that increased hyphal growth of *MAT***a** cells can also promote cell fusion.

**Figure 2.**
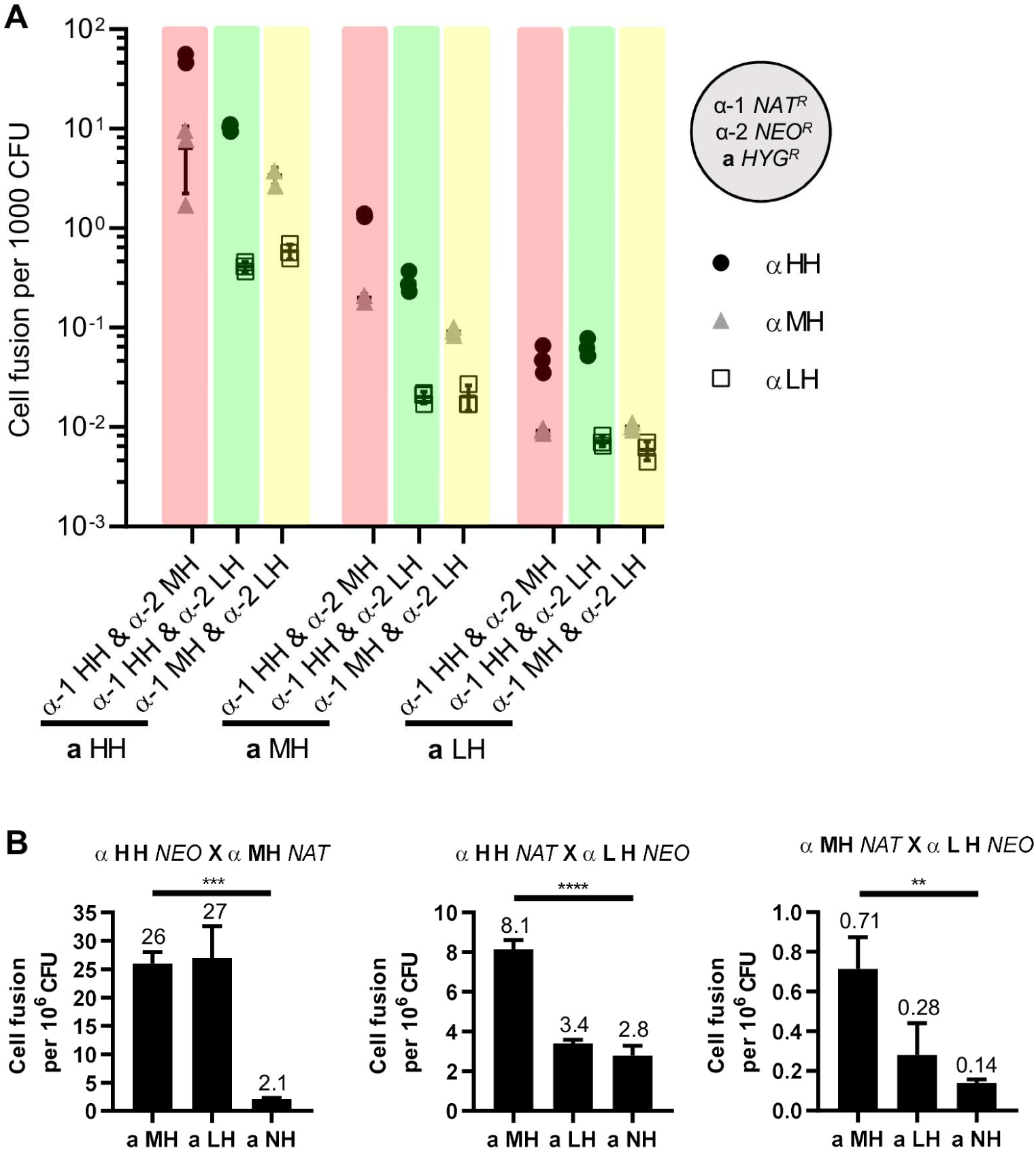
Unisexual reproduction potential enhances competition for mating partners of both mating types. (A) **a**-α Cell fusion frequencies for mating competitions between two *MAT*α strains over a *MAT***a** strain. (B) Unisexual α-α cell fusion frequencies in the presence of *MAT***a** strains of different hyphal growth phenotypes. ** indicates 0.001<*p*≤0.01, *** indicates 0.0001<p≤0.001, and **** indicates p≤0.0001 for each group analysis by one-way ANOVA.

Besides the observation that hyphal growth enhanced competition for mating partners of the opposite mating type, the presence of higher hyphal *MAT*α cells also stimulated **a**-α cell fusion. MH *MAT***a** and HH *MAT*α cells fused at a frequency of 52 cell fusion events per thousand CFU in the presence of MH *MAT*α cells compared to 10 cell fusion events per thousand CFU in the presence of LH *MAT*α cells (5.2-fold) (Dark yellow-shaded cells in Table S1). MH *MAT***a** and MH *MAT*α cells fused at a frequency of 6.4 cell fusion events per thousand CFU in the presence of HH *MAT*α cells compared to 3.4 cell fusion events per thousand CFU in the presence of LH *MAT*α cells (1.9-fold) (Dark blue-shaded cells in Table S1). Similar trends were observed during competition for LH *MAT***a** cells in that the presence of MH *MAT*α cells increased LH *MAT***a** and HH *MAT*α cell fusion frequency by 4.5-fold compared to the presence of LH *MAT*α cells (Medium yellow-shaded cells in Table S1), and the presence of HH *MAT*α cells increased LH *MAT***a** and MH *MAT*α cell fusion frequency by 2.2-fold compared to the presence of LH *MAT*α cells (Medium blue-shaded cells in Table S1). However, cell fusion frequencies between *MAT***a** cells and LH *MAT*α cells were comparable in the presence of HH or MH *MAT*α cells (0.41 or 0.59 cell fusion events per thousand CFU for MH *MAT***a** cells, and 0.02 or 0.02 cell fusion events per thousand CFU for LH *MAT***a** cells) (Dark and medium green-shaded cells in Table S1). Interestingly, no enhancement of cell fusion frequency by high hyphal *MAT*α cells was observed during competition for NH *MAT***a** cells (light color-shaded cells in Table S1). Notably, the enhancement of cell fusion frequency by higher hyphal *MAT*α cells did not occur when either *MAT***a** NH or *MAT*α LH cells were involved in **a**-α cell fusion, suggesting that strains with poor unisexual reproduction potential have a disadvantage in competing for mating partners of the opposite mating type.

The ability to undergo unisexual reproduction also correlated with cell fusion between cells of the same mating type when a *MAT***a** partner is present. In the presence of MH or LH *MAT***a** cells, HH and MH *MAT*α cells fused at higher frequencies (26 and 27 cell fusion events per million CFU, respectively) compared to HH and LH *MAT*α cells (8.1 and 3.4 cell fusion events per million CFU, respectively), and in the presence of MH, or LH, or NH *MAT***a** cells, HH and LH *MAT*α cells fused at higher frequencies (8.1, 3.4, and 2.8 cell fusion events per million CFU, respectively) compared to MH and LH *MAT*α cells (0.71, 0.28, and 0.14 cell fusion events per million CFU, respectively) (Figure 2B), suggesting that increased hyphal growth correlated with enhanced α-α cell fusion frequency in the presence of *MAT***a** cells. We also observed a trend where α-α cell fusion frequencies (HH and MH, HH and LH, and MH and LH) decreased with reduced hyphal *MAT***a** cells (Figure 2B), suggesting that the presence of more robust hyphal *MAT***a** cells can further enhance α-α cell fusion. In summary, strains with robust hyphal production have an advantage in competing for mating partners of the opposite mating type, and also for mating partners of the same mating type when cells of the opposite mating type are present.

### *gpa3*Δ mutation enhances competition for mating partners of the same or opposite mating type

The pheromone response pathway plays an important role in the yeast-to-hyphal morphological transition during *C. deneoformans* sexual reproduction. This signaling cascade is controlled by G proteins and RGS proteins, including the Gα protein Gpa3 which represses hyphal growth during mating [40-43]. To further examine the impact of the ability to undergo unisexual reproduction during mating competition, we generated strains enhanced for hyphal production by deleting the *GPA3* gene in the LH strain JEC21α. *gpa3*Δ mutants exhibited significantly increased hyphal growth during both unisexual and bisexual reproduction compared to the parental strain (Figure 3A). Next, mating competition assays were conducted using the enhanced hyphal (EH) strain JEC21α *gpa3*Δ to test its ability to compete for mating partners.

**Figure 3.**
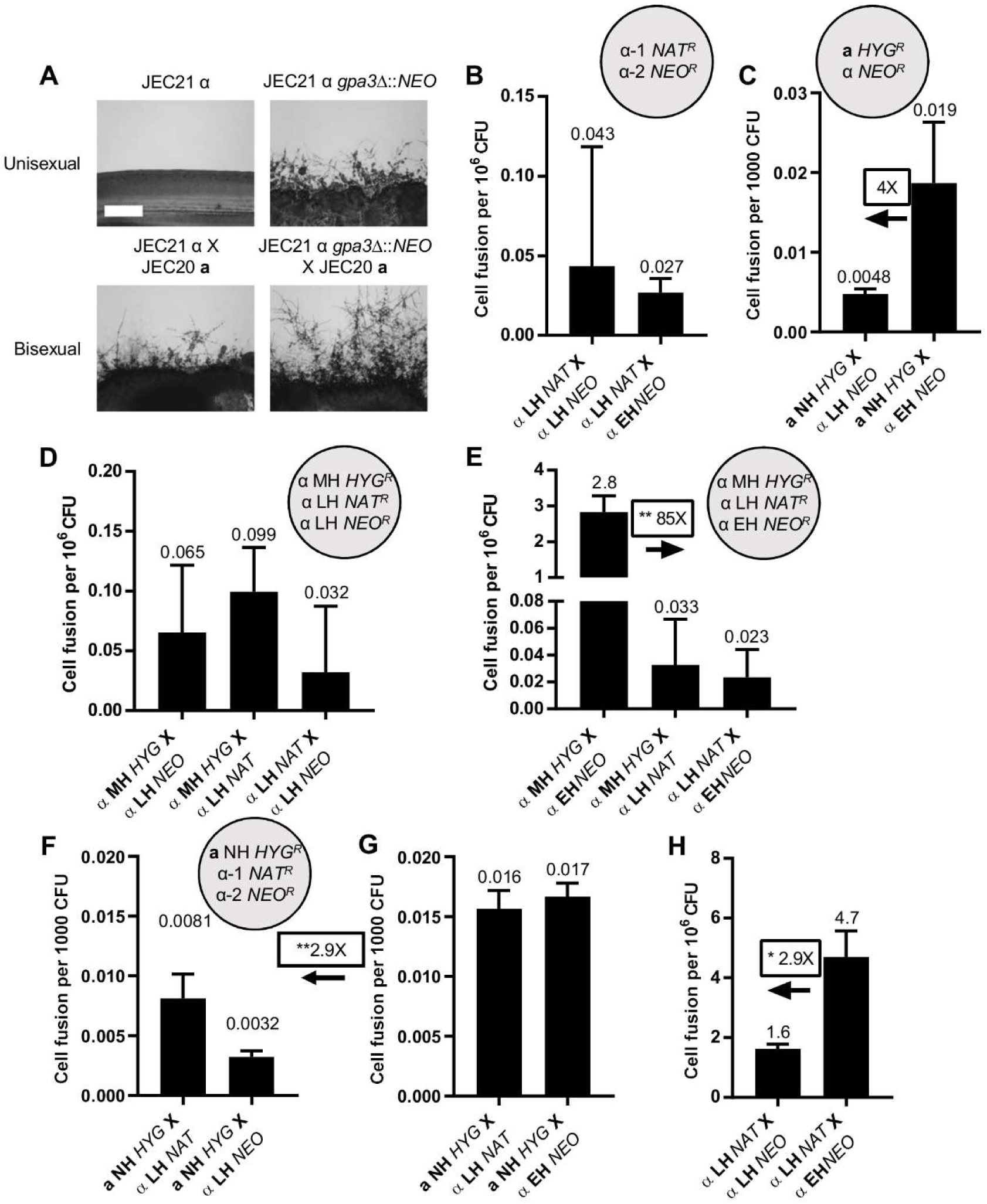
*gpa3*Δ mutation-enhances hyphal growth and competition for mating partners of both the same and opposite mating types. (A) Hyphal growth on V8 agar medium for one week for enhanced hyphal strain (JEC21α *gpa3*Δ::*NEO*) during unisexual (top) and unilateral bisexual (bottom) development compared to low hyphal strain (JEC21α). The scale bar represents 500 *µ*m. (B-C) Unisexual α-α (B) and bisexual **a**-α (C) cell fusion frequencies between two low hyphal strains and between no and enhanced hyphal strains. (D-E) Cell fusion frequencies during unisexual mating competition among two low and one intermediate hyphal *MAT*α strains (D), and one low, one enhanced, and one intermediate hyphal *MAT*α strains (E). (F-G) Cell fusion frequencies during bisexual mating competition for a no hyphal *MAT***a** strain between two low hyphal *MAT*α strains (F), and between one low and one enhanced hyphal *MAT*α strain (G). (H) Unisexual α-α cell fusion frequencies between two low hyphal *MAT*α strains, and between one low and one enhanced hyphal *MAT*α strain in the presence of a no hyphal *MAT***a** strain. * indicates 0.01<*p*≤0.05 for the pairwise comparison.

Similar to the observation in HH, MH, and LH strains, enhanced hyphal production did not increase cell fusion between *MAT*α cells but did increase cell fusion frequency by 4-fold between *MAT***a** and *MAT*α cells compared to the parental LH strain (Figure 3B, C). However, the increase is not statistically significant due to the low cell fusion frequencies between strains of low hyphal background. Unisexual mating competition assays were performed to compare the abilities of LH and EH *MAT*α cells to fuse with MH *MAT*α cells. In the control assay, cell fusion frequencies were comparable between cells of all three strain combinations (MH with LH-*NAT*, MH with LH-*NEO*, and LH-*NAT* with LH-*NEO*) (Figure 3D). In the assay mixing LH, MH, and EH cells, EH cells fused with MH cells at a significantly higher frequency of 2.8 cell fusion events per million CFU compared to LH cells (85-fold) (Figure 3E), suggesting that deletion of *GPA3* increases competitiveness for mating partners of the same mating type. In the mating competition during bisexual reproduction, no advantage was observed in the fusion of NH *MAT***a** cells with either EH or the parental LH *MAT*α cells (Figure 3G). However, a significant 2.9-fold increase was observed in total **a**-α cell fusion events during mating competitions for NH *MAT***a** cells between EH and LH *MAT*α cells compared to control mating competitions for the same *MAT***a** cells between LH-*NAT* and LH-*NEO MAT*α cells (Figure 3F-3G), indicating that presence of cells with enhanced ability to undergo unisexual reproduction allows both EH and LH *MATa* cells to fuse with *MAT***a** mating partners more efficiently during bisexual reproduction. A significant 2.9-fold increase was observed in α-α cell fusion between EH and LH *MAT*α cells in the presence of NH *MAT***a** cells compared to cell fusion between LH-*NAT* and LH-*NEO MAT*α cells (Figure 3H), suggesting that in the presence of *MAT***a** cells, *GPA3* deletion also enhances competition for mating partners of the same mating type. Overall, this analysis of the enhanced hyphal growth strain JEC21α *gpa3*Δ provides additional support for models in which increased unisexual reproduction potential enhances competition for mating partners.

### Hyphal growth promotes foraging for mating partners

Unisexual reproduction provides evolutionary and ecological benefits for *C. deneoformans* by generating aneuploid progeny with phenotypic diversity and by promoting habitat exploration through hyphal growth [29, 34]. Here we further show that unisexual cells have an advantage in competing for mating partners within the same colony. We tested whether hyphal growth during unisexual reproduction confers benefits in foraging for mating partners.

Both long-term and short-term foraging for mating partner experiments suggested that hyphal growth promoted foraging for mating. In a six-week mating confrontation experiment, hyphae of different *MAT*α unisexual reproduction backgrounds marked with NEO grew towards the same *MAT***a** cells marked with HYG that were 4 mm apart (Figure 4A, B). When competing for either the same *MAT***a** or *MAT*α cells (except LH *MAT*α cells), although not all pairwise comparisons by t-test were significant due to the lack of contact when strains of no or low hyphal growth were involved, a significant trend by one-way ANOVA was observed that isolates with more hyphal growth yielded more double drug resistant colonies than isolates with reduced hyphal growth (Figure 4C and Table S2). In a seven-day mini-colony mating experiment, colonies derived from single cells produced hyphae that allowed contact with adjacent colonies of the opposite mating type (Figure S2A). Similar to the long-term mating confrontation experiment, a significant trend by one-way ANOVA was observed in which isolates with more hyphal growth had an advantage in forming double drug resistant colonies (Figure S2B and Table S3). Although pairwise comparisons by t-test showed significant differences between crosses involved NH and LH cells that were not observed in the confrontation experiment, this discrepancy is due to differences in the experimental setup where cells were inoculated at 0.4 cm apart during confrontation, whereas randomly plated on agar media during mini-colony mating experiment, where colony contact is enabled by both hyphal growth and chance. Overall, these results suggest that hyphal growth during unisexual reproduction can facilitate contact between mating partners in adjacent environments.

**Figure 4.**
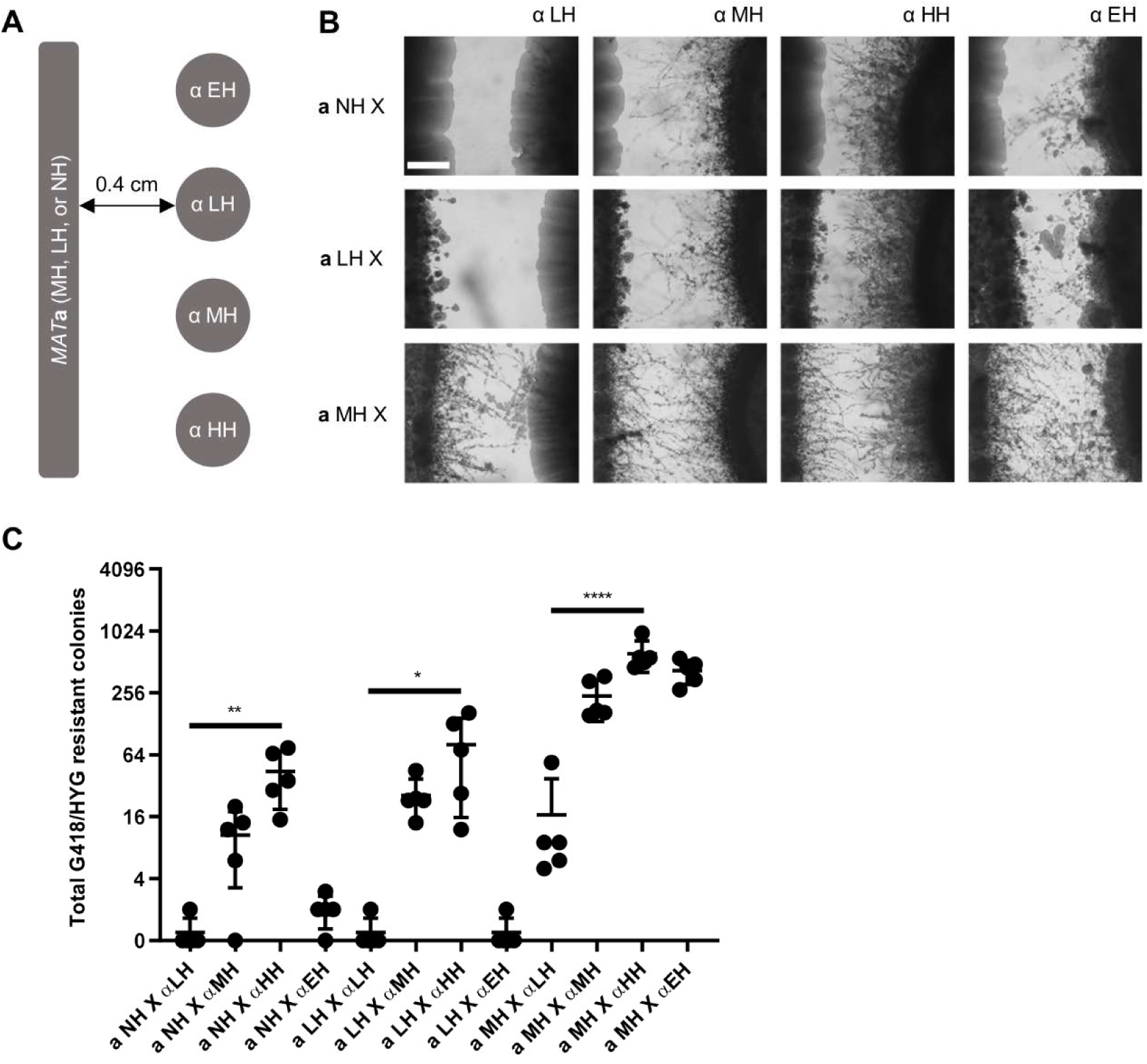
Hyphal growth during unisexual reproduction promotes foraging for mating partners in a mating confrontation assay. (A) Schematic diagram for the confrontation mating experiment setup. *MAT***a** cells were grown to form a line of cells, and *MAT*α cells were spot-inoculated 4 mm apart in parallel along the *MAT***a** cells. (B) LH, MH, HH, and EH *MAT*α hyphal cells were grown towards NH, LH, and MH *MAT***a** cells for six weeks. The scale bar represents 500 *µ*m. (C) Total G418/HYG resistant colonies produced by each confrontation mating pair. LH, MH, HH, and EH *MAT*α hyphal cells yielded an average of 0.4, 10, 44, and 1.8 double drug resistant colonies with NH *MAT***a** cells, 0.4, 26, 81, and 0.4 double drug resistant colonies with LH *MAT***a** cells, 17, 239, 613, and 422 double drug resistant colonies with MH *MAT***a** cells, respectively. * indicates 0.01<*p*≤0.05, ** indicates 0.001<p≤0.01, and **** indicates p≤0.0001 for each group analysis by one-way ANOVA.

### Pheromone response pathway activation is correlated with hyphal growth phenotype during bisexual reproduction

Elevated pheromone response pathway activation and increased response to pheromones are critical to successful courtship during mating in *S. cerevisiae* and *C. albicans* [10, 11, 14]. *S. cerevisiae* utilizes the α-factor protease Bar1 and **a**-factor barrier Afb1 to discriminate mating partners with different pheromone levels and drive evolution towards higher pheromone production for efficient mating [44-47]. Pheromones also stimulate mating and the yeast-to-hyphal morphological transition during *C. neoformans* bisexual reproduction [48]. To determine the role of the pheromone response pathway during *C. deneoformans* mating competition, expression levels were examined for the genes encoding the pheromones MFα and MF**a**, the pheromone receptors Ste3α and Ste3**a**, the MAP kinase Cpk1, the transcription factors Mat2 and Znf2, and the plasma membrane fusion protein Prm1 [23, 40, 49].

Pheromone response pathway activation did not correlate with the hyphal growth phenotype in *MAT***a** or *MAT*α strains. In *MAT***a** strains, all of the pheromone response pathway genes were significantly upregulated in the MH strain XL280**a** compared to the LH and NH strains, but only *MF***a** and *PRM1* were significantly upregulated in the LH strain XL187**a** compared to the NH strain JEC20**a** (Figure S3). In *MAT*α strains, all of the pheromone response pathway genes were significantly upregulated in the HH and MH strains compared to the LH strain JEC21α, but the MH strain XL280α had a significantly higher pheromone pathway activation compared to the HH strain XL190α (Figure S3). The pheromone pathway was significantly upregulated in the EH strain JEC21α *gpa3*Δ compared to the parental LH strain JEC21α (Figure S3). These expression analyses suggest that pheromone response pathway activation *per se* is not sufficient to explain the ability to undergo unisexual reproduction and its association with competition for mating partners. It was previously shown that the cell fusion protein Prm1 is not required for unisexual reproduction [23], and certain environmental factors, such as copper and glucosamine, can induce hyphal growth independently of the pheromone response pathway [20, 21]. In a recent study on the quorum sensing peptide Qsp1, deletion of pheromone and pheromone receptor genes had little impact on hyphal growth during unisexual reproduction [22], further indicating the polygenic nature of unisexual reproduction.

Despite the incongruent association of the pheromone response pathway and the ability to undergo unisexual reproduction, pheromone response pathway activation was positively correlated with the hyphal growth during bisexual reproduction. The α pheromone gene *MF*α, both **a** and α pheromone receptor genes *STE3*α and *STE3***a**, and the plasma membrane fusion gene *PRM1* showed significant correlation with the hyphal growth phenotype (Figure 5A). Although *MF***a** expression was lower in the cross between **a** MH and α HH strains compared to **a** LH and α MH strains, the significant upregulation of the *MF*α, *STE3*α, and *STE3***a** may compensate for the overall pheromone response activation (Figure 5A). The gene expression patterns of two transcription factors Mat2 and Znf2 that regulate yeast-to-hyphal morphological transition and mating significantly correlated with the hyphal growth except for the cross between **a** MH and α HH strains. Nonetheless, these two genes were expressed at higher levels compared to the cross between **a** LH and α MH strains (Figure 5A). The expression of the MAP Kinase *CPK1* gene poorly correlated with hyphal growth (Figure 5A), which is likely due to post-translational control of the MAP kinase through phosphorylation, which can relieve a requirement for expression level upregulation during pathway activation. Overall, the pheromone response pathway activation is largely congruent with the hyphal growth phenotype suggesting that in the presence of cells of the opposite mating type, unisexual cells are capable of upregulating the pheromone response pathway in both *MAT***a** and *MAT*α cells to compete for mating partners.

**Figure 5.**
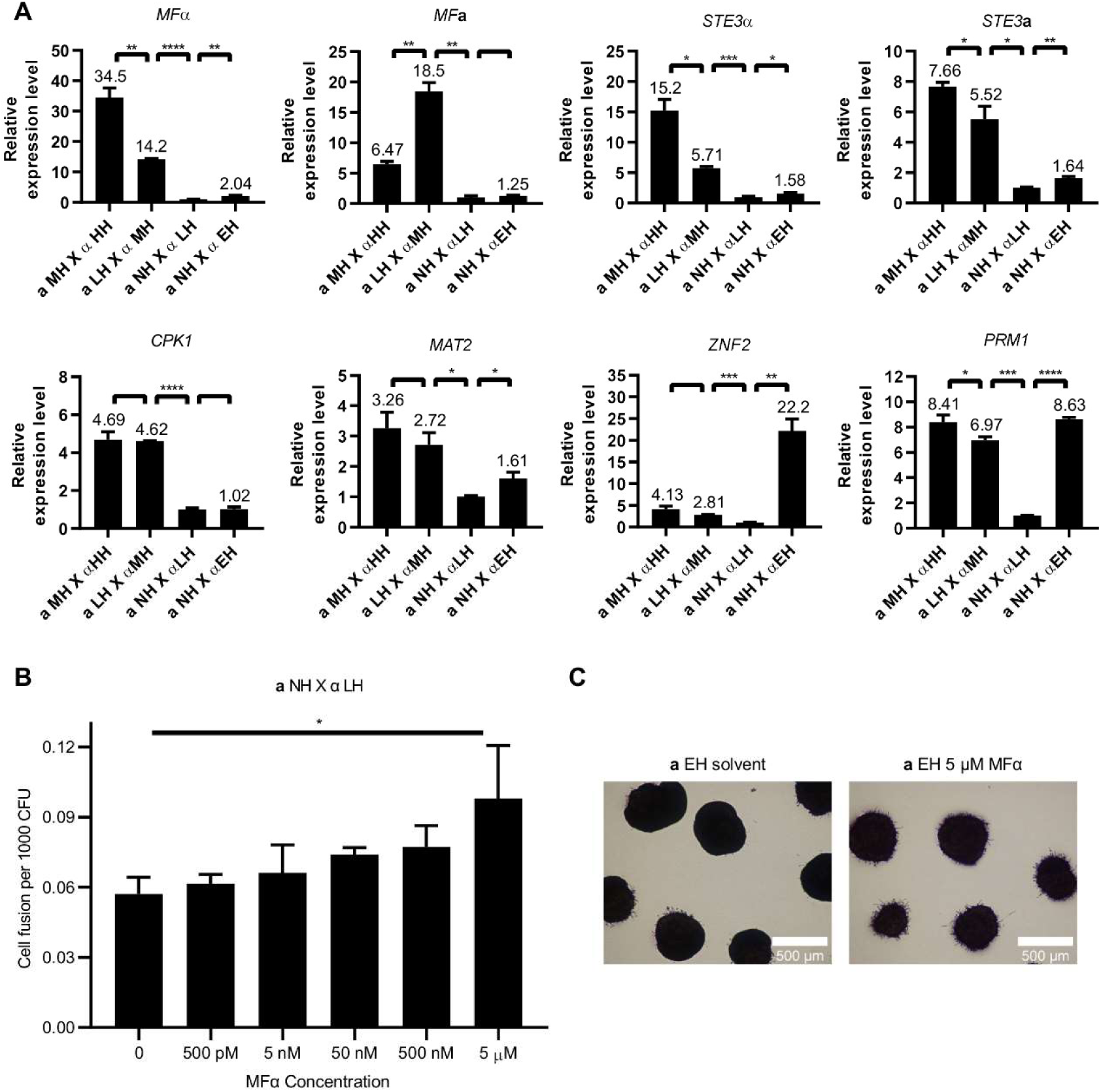
Pheromone response pathway activation associated with unisexual reproduction. (A) Pheromone response pathway activation during bisexual reproduction is correlated with the hyphal growth phenotype. Gene expression patterns for *MF*α, *MF***a**, *STE3*α, *STE3***a**, *CPK1, MAT2, ZNF2*, and *PRM1* were examined by qRT PCR (NS indicates p≤0.01, * indicates 0.01<*p*≤0.05, ** indicates 0.001<*p*≤0.01, *** indicates 0.0001<p≤0.001, and **** indicates p≤0.0001 for each pairwise comparison.). Crosses between MH *MAT***a** (XL280**a**) and HH *MAT*α (XL190α) strains, LH *MAT***a** (XL187**a**) and MH*MAT*α (XL280α) strains, NH *MAT***a** (JEC20**a**) and LH *MAT*α (JEC21α) strains, and NH *MAT***a** (JEC20**a**) and EH *MAT*α (JEC21α *gpa3*Δ::*NEO*) strains were grown on V8 agar medium for 36 hours. The expression levels of genes from the cross between JEC20**a** and JEC21α were set to 1, and the remaining values were normalized to this. The error bars represent the standard deviation of the mean for three biological replicates. (B) Exogenous pheromone enhances cell fusion frequency between NH *MAT***a** and LH *MAT*α strains in a dose dependent manner. *p*=0.0125 by one-way ANOVA. (C) EH *MAT***a** (JEC20**a** *gpa3*Δ::*ADE2 ade2*) colonies produced more hyphae in response to exogenous pheromone after three-day incubation on MS medium.

To validate that pheromone contributes to mating competitiveness, we tested whether synthetic α pheromone promotes **a**-α cell fusion. Indeed, exogenous α pheromone promoted cell fusion between **a** NH and α LH cells in a dose dependent manner (Figure 5B). However, the enhancement of cell fusion frequency by pheromone is limited, compared to the 2788-fold increase of cell fusion frequency between **a** MH and α HH cells over **a** NH and α LH cells (Figure 1D) that coincided with a 34.5-fold increase in α pheromone expression (Figure 5A). Interestingly, the mild increase in cell fusion by exogenous pheromone is not observed in the cross between **a** NH and α EH cells (Figure S4A), which is likely due to the saturation of Ste3**a** by the 1184-fold higher α pheromone expressed by α EH cells (Figure S3). The less than two-fold increase in cell fusion provided by 5 μM exogenous pheromone suggested that changes in α pheromone alone are not able to tip the balance and drive the entire pheromone response pathway towards a stronger increase in mating efficiency. In support, exogenous supply of 500 nM pheromone provided limited impact in cell fusion between cells of higher hyphal growth phenotypes (Figure S4A); and when hyphal growth was suppressed under nutrient rich conditions, exogenous pheromone had little to no impact on cell fusion (Figure S4B).

In response to the pheromone signal, *S. cerevisiae* undergoes filamentous growth to enhance the probabilities of cells finding a mating partner [50]. Here we observed that the **a** EH colonies responded to 5 μM α pheromone peptide and produced abundant hyphae (Figure 5C), similar to previous report [51], suggesting that pheromone can promote hyphal growth and increase the contact opportunities between adjacent colonies within the same environment.

### *gpa3*Δ mutation resulted in a fitness cost

Upregulation of the pheromone response pathway enhances mating efficiency; however, this upregulation can result in a fitness cost in *S. cerevisiae* and *C. albicans* [14, 52]. In yeast, a short-term experimental evolution experiment showed that mutations abrogating expression of 23 genes involved in mating conferred a fitness benefit during yeast growth when functions of these genes are not required [52]. In *C. albicans*, cells undergo a white-opaque switch and upregulate the pheromone response pathway, which results in a fitness cost for the opaque cells [14]. Given that the pheromone response pathway is activated at a higher level in *C. deneoformans* strains with more hyphal growth during bisexual reproduction (Figure 5A), we investigated whether unisexual reproduction confers a fitness cost.

Growth curve analyses in YPD liquid medium were performed using an automated Tecan Sunrise absorbance reader to determine the fitness of different strains. The cultures were agitated with vigorous shaking for one minute bihourly before each OD_600_ measurement. The minimum agitation allows for differentiation of growth curve kinetics among the strains tested, which would otherwise be indistinguishable when grown on solid YPD medium. Compared to the low hyphal growth strains JEC21α and JEC20**a**, *gpa3*Δ mutants exhibited a growth defect in nutrient rich media, suggesting that deletion of *GPA3* resulted in a fitness cost (Figure S5). However, this fitness cost was not due to the yeast-to-hyphal morphological transition as hyphal growth was not present. Interestingly, deletion of the *GPA3* gene in the sister species *C. neoformans* did not induce hyphal growth or cause a growth defect compared to the non-hyphal strain KN99**a** (Figure S5). It has been shown that the pheromone response pathway activation by *GPA3* gene deletion is lower in KN99**a** than in JEC21 or JEC20, indicating that deletion of *GPA3* is not sufficient to rewire cellular responses to induce unisexual reproduction in *C. neoformans* [40].

We next performed fitness competition experiments. After 10 days of incubation of equally mixed cells on YPD and V8 agar medium, cells were collected and plated on selection media to determine colony forming units for each competition strain. Hyphal growth was observed on both YPD and V8 media when **a** MH cells were present, and both yeast cells and hyphae were collected to compare fitness (Figure S6). During competition between two LH strains, cells were recovered at about 1 to 1 ratio after 10-day incubation on both YPD and V8 media both in the absence or in the presence of **a** cells (Figure 6A). In contrast, LH strain outcompeted EH strain when cells were incubated on V8 medium or when **a** cells were present (Figure 6B). On YPD medium in the absence of **a** cells, the EH strain that displayed poor growth in liquid media (Figure S5) outcompeted the parental LH strain(Figure 6B), which is likely due to differential cellular responses under different growth conditions. However, this competition advantage was reversed when **a** cells were present or when mixed cells were incubated on V8 medium (Figure 6B), suggesting that the presence cells of the opposite mating type or the mating inducing environment can elicit a fitness cost. This competition disadvantage for the EH strain on V8 medium was further exacerbated when **a** cells were present during competition (Figure 6B). These fitness competition assays demonstrate that *gpa3*Δ mutation enhances mating competition at a cost of growth fitness, and this fitness cost is likely due to the energy required in the expression of the pheromone response pathway genes. It has been reported that long-term passage on rich media in the lab often diminishes the hyphal growth phenotype of *MAT***a** strains, which further suggests that there is a fitness cost associated with the ability to undergo unisexual reproduction [53]. Interestingly, in *S. cerevisiae*, the [*SWI*^+^] prion state promotes outcrossing efficiency due to a defect in *HO* expression and mating type switching, which also leads to a fitness cost [54]. Taken together, fungal species have evolved different strategies in promoting mating in nature accompanied with a fitness tradeoff.

**Figure 6.**
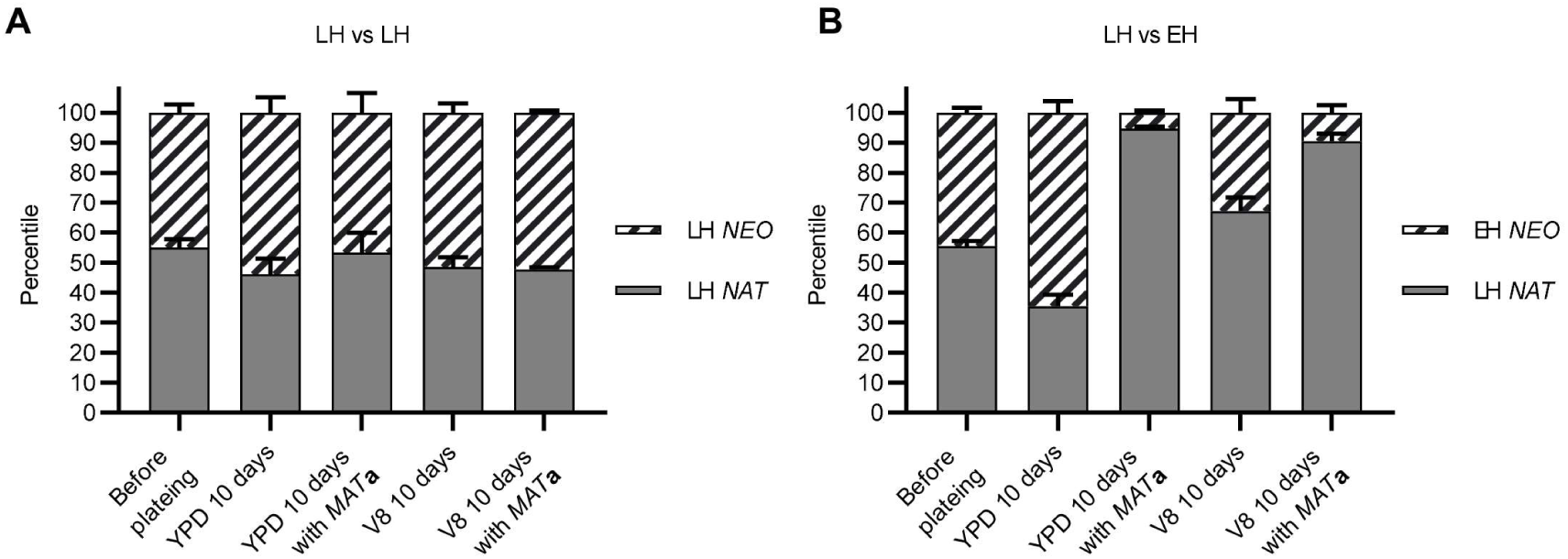
*gpa3*Δ mutation confers a fitness cost under mating inducing conditions. Equal numbers of cells of (A) two low hyphal strains (JEC21α *NAT* and JEC21α *NEO*) and of (B) a low hyphal strain (JEC21α *NAT*) and an enhanced hyphal strain (JEC21α *gpa3*Δ::*NEO*) were co-cultured on YPD and V8 agar medium in the absence or the presence of equal number of cells of an intermediate hyphal **a** strain (XL280**a**). CFUs of each strain were counted before plating and after 10 days of incubation to determine competition fitness.

## Conclusion

Sexual reproduction plays a pivotal role in shaping fungal population structure and diversity in nature. However, studies on how fungi secure a mating partner in nature for successful mating are limited. In this study, we aimed to characterize the ecological and evolutionary benefits of unisexual reproduction in *C. deneoformans*. Similar to the landmark study by Jackson and Hartwell showing that higher pheromone production promotes courtship in *S. cerevisiae* [11], we showed that strains with higher potential for unisexual reproduction are more competitive for mating partners of both the same and the opposite mating types when cells of the opposite mating type are present, and the pheromone response pathway activation is positively correlated with the hyphal growth phenotype (Figure 7). More interestingly, in addition to pheromone sensing, unisexual cells employ hyphal growth to increase contact opportunities between colonies at relatively long distances. However, this mating competition advantage results in a fitness cost for unisexual cells during mitotic growth under mating-inducing conditions.

**Figure 7.**
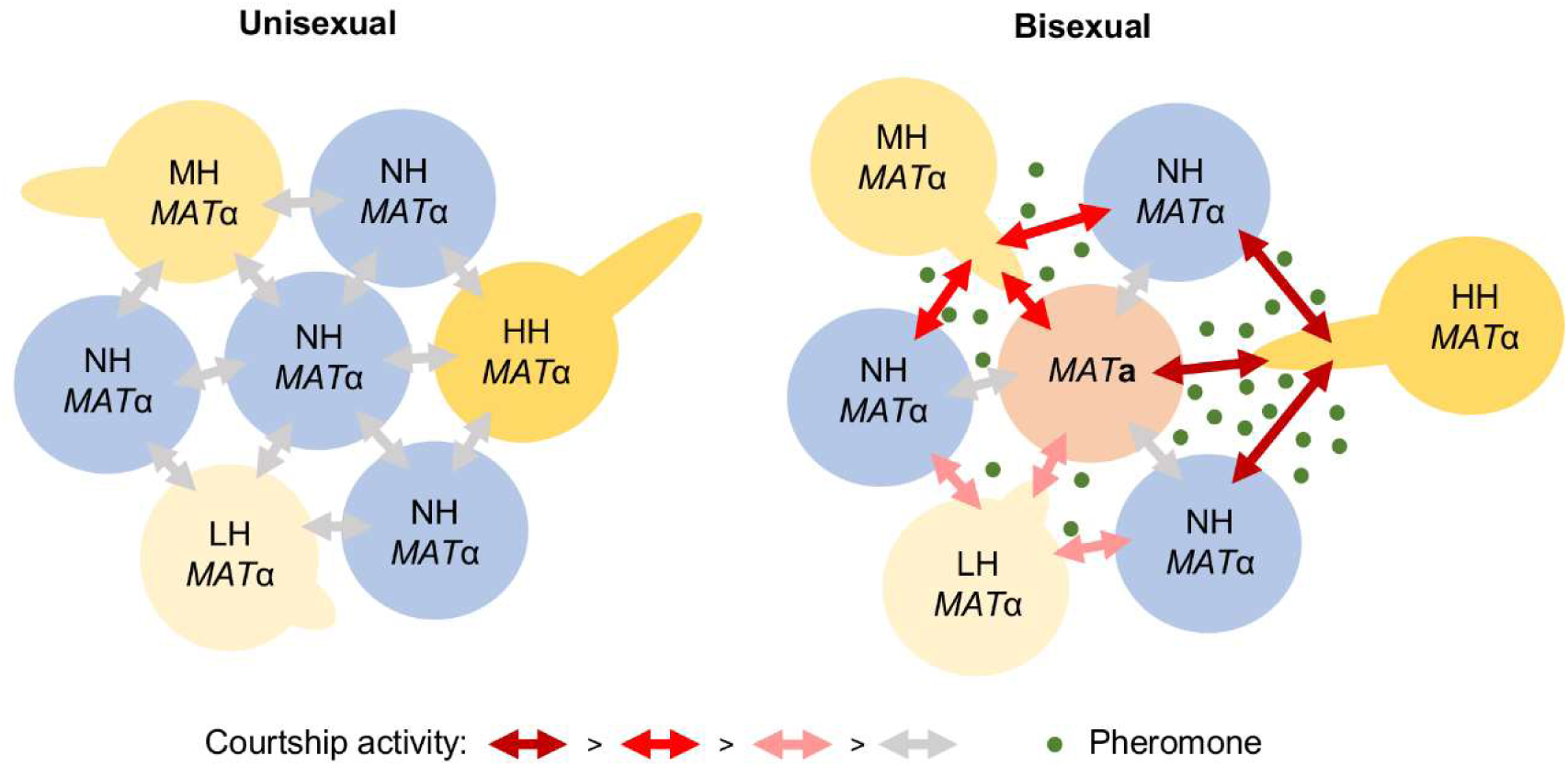
Model for the role of unisexual reproduction in foraging for mating partners. In the absence of the opposite mating type, the ability to undergo unisexual reproduction does not promote foraging for mating partners of the same mating type (left). In the presence of the opposite mating type, hyphal growth promotes foraging for mating partners of both the same and the opposite mating type (right). NH: no hyphal growth; LH: low hyphal growth; MH: intermediate hyphal growth; HH: high hyphal growth.

The strains involved in this study were all derived from natural and clinical isolates under laboratory conditions, suggesting that the ability to undergo unisexual reproduction is likely to span a broad range in the environment. The majority of natural and clinical isolates of the *Cryptococcus* species complex are found to be of the α mating type, which accounts for 99% of *C. neoformans* isolates [25-27]. In a survey of *C. deneoformans* environmental distribution around the Mediterranean basin, 27% of isolates are *MAT***a** isolates, which were all recovered in Greece, suggesting certain environmental niches harbor *MAT***a** cells [27]. Genomic and genetic evidence also suggest that recombination is prevalent among these environmental isolates, and some isolates are isolated from a single *Eucalytus* hollow, which underscores that mating occurs in nature [30-32]. Although sexual structures of *Cryptococcus* species have yet to be documented in nature, plant material-based media such as V8, on which we conducted the mating competition assays, can readily induce sexual reproduction under laboratory conditions, suggesting that the mating competition we observed could happen in its environmental niche. We hypothesize that in the presence of *MAT***a** cells sparsely distributed in the environment, undirected hyphal growth first enables unisexual *MAT*α cells to forage for mating partners over a much larger surface area than is available to cells within a much more compact budding yeast colony. Next, as *MAT*α hyphae come into the proximity of rare *MAT***a** cells, pheromone response pathway activation in both *MAT*α and *MAT***a** cells can further enhance mating competition. This mating competition advantage could promote outcrossing and provide an evolutionary advantage by facilitating genome reshuffling via meiotic recombination in a pathogenic yeast species.

## Materials and methods

### Strains, media, and growth conditions

Strains used in this study are listed in Table S4. Strains with different hyphal growth phenotypes, XL190α, XL280α, JEC21α, XL280**a**, XL187**a**, and JEC20**a**, were selected to represent high, intermediate, and low hyphal strains [16, 35, 38, 39]. Yeast cells were grown at 30°C on Yeast extract Peptone Dextrose (YPD) medium. Strains harboring dominant selectable markers were grown on YPD medium supplemented with 100 μg/mL nourseothricin (NAT), 200 μg/mL G418 (NEO), or 200 μg/mL hygromycin (HYG) for selection. Mating assays were performed on 5% V8 juice agar medium (pH = 7.0) or Murashige and Skoog (MS) medium minus sucrose (Sigma-Aldrich) in the dark at room temperature for the designated time period.

### Drug-resistant marker strain generation and gene deletion

NAT (pAI3) or G418/NEO (pJAF1) resistant expression constructs were introduced into XL190α, XL280α, and JEC21α, and a HYG (pJAF15) resistant expression construct was introduced into XL280α, XL280**a**, XL187**a**, and JEC20**a** ectopically via biolistic transformation as previously described [55-57].

To generate deletion mutants for *GPA3*, a deletion construct consisting of the 5’ upstream and 3’ downstream regions of *GPA3* gene flanking the *NEO* cassette was generated by overlap PCR as previously described [58]. The *GPA3* deletion construct was introduced into the strain JEC21α via biolistic transformation. Transformants were selected on YPD medium supplemented with G418, and gene replacement by homologous recombination was confirmed by PCR. Primers used to generate these deletion constructs are listed in Table S5.

### Microscopy

Cells were grown on V8 agar medium for seven days or three weeks in the dark at room temperature to allow hyphal formation. Hyphal growth on the edge of mating patches was imaged using a Nikon Eclipse E400 microscope equipped with a Nikon DXM1200F camera.

### Competitive mating assays

For each competitive mating assay, cells were grown overnight in YPD liquid medium at 30°C and adjusted to OD_600_=0.5 in sterile H_2_O, and then equal volumes of cells marked with different dominant drug resistant markers were mixed and spot inoculated (50 μl) on V8 agar medium. The mating plates were incubated for four days in the dark at room temperature. The cells were harvested and plated in serial dilution on YPD medium and YPD medium supplemented with different two drug combinations (NAT and NEO, NAT and HYG, or NEO and HYG). The cells were incubated for three to five days at room temperature and colony forming units were counted. Cell fusion frequencies were determined as double drug resistant CFU/total CFU. The complete competitive mating experimental design is listed in Table S6. Each mating competition was performed in biological triplicate.

### Foraging for mating assays

To investigate whether hyphal growth enables cells foraging for mating partners, we performed long-term confrontation and short-term mini-colony mating experiments. For the confrontation mating experiment, HYG resistant XL280**a**, XL187**a**, and JEC20**a** were streaked and grown on V8 medium to form a line of cells, and then NEO resistant XL190α, XL280α, and JEC21α, and JEC21α *gpa3*Δ::*NEO* were spot inoculated 4 mm apart in parallel along the *MAT***a** cells. Unisexual hyphae grew towards cells of the opposite mating type for six weeks. Then the cells were collected and plated on YPD medium supplemented with HYG and NEO. After incubation at room temperature for three to five days, total double drug resistant colony forming units were counted to determine whether hyphal growth conferred an advantage in foraging for mating partners. Each confrontation mating experiment was performed in biological quintuplicate. For the mini-colony mating experiment, the aforementioned HYG resistant *MAT***a** strains and NEO resistant *MAT*α strains were grown overnight in YPD liquid medium and adjusted to OD_600_=0.008. For each mating pair, 100 μl of *MAT***a** cells and 100 μl of *MAT*α cells were mixed and plated on V8 agar medium to form evenly spaced mini colonies. Unisexual hyphae facilitate contact between adjacent colonies after growing for seven days. Then the cells were collected and plated on YPD medium and YPD medium supplemented with HYG and NEO. The cells were incubated for three to five days at room temperature and colony forming units were counted. Cell fusion frequencies were determined as double drug resistant CFU/total CFU. Each mating was performed in biological triplicate.

### RNA extraction and qRT-PCR

To examine pheromone response pathway activation, qRT PCR experiments were performed on RNA extracted from cells incubated on V8 agar medium for 36 hours as previously described [23]. In brief, XL190α, XL280α, JEC21α, JEC21α *gpa3*Δ::*NEO*, XL280**a**, XL187**a**, and JEC20**a** were grown overnight in YPD liquid medium and adjusted to OD_600_=2 in sterile H_2_O. Then cells of individual strains and an equal-volume mixtures of cells for crosses between XL190α and XL280**a**, XL280α and XL187**a**, JEC21α and JEC20**a**, and JEC21α *gpa3*Δ::*NEO* and JEC20**a** were spotted (250 μl) on V8 medium and incubated for 36 hours. Cell patches of individual strains and of mixture of **a** and α strains were scraped off the medium and transferred into Eppendorf tubes then flash frozen in liquid nitrogen. RNA was extracted using TRIzol reagent (Thermo) following the manufacturer’s instructions. RNA was treated with Turbo DNAse (Ambion), and single-stranded cDNA was synthesized by AffinityScript RT-RNAse (Stratagene). cDNA synthesized without the RT/RNAse block enzyme mixture was used to control for genomic DNA contamination. The relative expression levels of targeted genes were measured by qRT PCR using Brilliant III ultra-fast SYBR green QPCR mix (Stratagene) in an Applied Biosystems 7500 Real-Time PCR system. A “no template control” was used to analyze the resulting melting curves to exclude primer artifacts for each target gene. Gene expression levels were normalized to the endogenous reference gene *GPD1* using the comparative ΔΔCt method. Primers used for qRT-PCR are listed in Table S5. For each target gene and each sample, technical triplicate and biological triplicate were performed.

### Mating assays with exogenous α pheromone peptide

To address whether pheromone promotes mating competition and hyphal growth, carboxyl farnesylated and methylated α pheromone peptide (QEAHPGGMTLC) (synthesized at GenScript, USA) was tested for its impact on mating and hyphal growth. α pheromone peptide was dissolved in methanol at the concentration of 50 μM and 10-fold serial dilutions in methanol were prepared as stock solutions. For the mating assay, *HYG* marked *MAT***a** and *NEO* marked *MAT*α cells were prepared and mixed as mentioned above for the mating competition assays. α pheromone peptide was added to mixed **a** NH (CF926) and α LH (CF759) cells at the concentrations of 0, 500 pM, 5 nM, 50 nM, 500 nM, and 5 μM, and the mixed cells were spot-inoculated on the V8 media and incubated in the dark at room temperature for four days. Cells were then harvested and plated on both YPD and YPD media supplemented with NEO and HYG to determine cell fusion frequency. Same mating assays were carried out for crosses between **a** MH (CF978) and α HH (CF914), **a** LH (CF931) and α MH (CF752), **a** NH (CF926) and α LH (CF759), and **a** NH (CF926) and α EH (CF1314) both in the absence and in the presence of 500 nM α pheromone peptide on both YPD and V8 media.

To test the impact of α pheromone peptide on hyphal growth, 5 μM α pheromone peptide and methanol were dropped onto MS media and allowed to dry. **a** EH (YPH86) cells were grown overnight and washed with H_2_O twice, and then inoculated onto the MS plate with dried α pheromone peptide and methanol droplets at a different spot. Cells were then microscopically manipulated and transferred to the α pheromone peptide and methanol spots. Colony hyphal growth was monitored daily and imaged after incubation in the dark at room temperature for 72 hours.

### Fitness competition and growth curve assays

Competition experiments were performed to compare fitness between two low hyphal strains and between low and enhanced hyphal strains both in the absence and in the presence of the **a** MH cells. Overnight cultures of JEC21α *NAT*, JEC21α *NEO*, JEC21α *gpa3*Δ::*NEO*, and XL280**a** were washed with H_2_O twice and cell densities were determined with a hemocytometer. For each competition experiment, 10 μl of H_2_O containing 100,000 cells of each strain was spotted on either YPD or V8 agar medium and incubated in the dark at room temperature for 10 days. Cells were collected and plated on YPD medium supplemented with NAT or G418 to determine colony forming units. Cell mixtures were plated before incubation to control for equal mixing. Each competition was performed in triplicate. Fitness was determined by calculating the percentile of the recovered CFU of each strain out of the total recovered dominant drug resistant CFU.

To determine the growth fitness of different unisexual strains, KN99**a**, KN99**a** *gpa3*Δ::*NEO*, JEC20**a**, JEC20**a** *gpa3*Δ::*NEO*, JEC21α, and JEC21α *gpa3*Δ::*NEO* were grown overnight in YPD liquid medium and washed twice in H_2_O. 10,000 cells for each strain were resuspended in 200 μl YPD liquid medium and incubated in a 96-well plate (Corning) at 30°C with vigorous shaking for 1 min bihourly. OD_600_ readings were measured bi-hourly after shaking using an automated Tecan Sunrise absorbance reader. Each sample was tested in quintuplicate.

### Statistical analysis

All statistical analyses were performed using the Graphpad Prism 7 program. Welch’s t-test was performed for each pairwise comparison, and one-way ANOVA was performed for each group analysis with a *p* value lower than 0.05 considered statistically significant (* indicates 0.01<*p*≤0.05, ** indicates 0.001<*p*≤0.01, *** indicates 0.0001<p≤0.001, and **** indicates p≤0.0001).

## Acknowledgements

We thank Shelby Priest, Daniel Lew, Sheng Sun, and Zanetta Chang for critical reading of the manuscript. This work was supported by NIH/NIAID R37 grant AI39115-21 and R01 grant AI50113-15 to J.H.

## Author Contributions

Conceptualization, C.F., T.P.T., and J.H.; Formal analysis, C.F., T.P.T., and J.H.; Funding acquisition, J.H.; Investigation, C.F. and T.P.T.; Methodology, C.F., T.P.T., and J.H.; Resources, C.F. and J.H.; Supervision, J.H.; Writing – original draft, C.F.; Writing – review & editing, C.F., T.P.T., and J.H.

## Supporting information

**Figure S1. Derivation of the strains used in this study**.

B4478 *MAT*α, JEC20**a**, XL187**a**, XL190α, and XL280α are F2 progeny from the cross between F1 progeny B3502 *MAT***a** and B3501 *MAT*α, which were derived from a cross between the environmental isolate NIH433 *MAT***a** and the clinical isolate NIH12 *MAT*α. JEC20**a** was then crossed with B4478 *MAT*α, and an α progeny was backcrossed with JEC20**a**. This process was repeated 9 times to yield the congenic partner B4500 JEC21α of B4476 JEC20**a**. XL280α was crossed with JEC20**a**, and a *MAT***a** progeny was backcrossed with XL280α. This process was repeated 9 times to yield the congenic partner XL280**a**.

**Figure S2. Hyphal growth promotes foraging for mating partners in a mini-colony mating assay**.

(A) LH, MH, HH, NH, and EH *MAT*α colonies derived from single cells were grown for seven days and in some cases, hyphae facilitated contact between *MAT*α and *MAT***a** colonies. The scale bar represents 500 *µ*m. (B) Cell fusion frequencies between *MAT***a** and *MAT*α mating partners for each mating pair are shown. * indicates 0.01<*p*≤0.05 and ** indicates 0.001<p≤0.01 for each group analysis by one-way ANOVA.

**Figure S3. Pheromone response pathway activation is not correlated with the hyphal growth phenotype**.

Gene expression patterns for *MF*α, *MF***a**, *STE3*α, *STE3***a**, *CPK1, MAT2, ZNF2*, and *PRM1* were examined by qRT PCR (NS indicates p≤0.01, * indicates 0.01<*p*≤0.05, ** indicates 0.001<*p*≤0.01, *** indicates 0.0001<p≤0.001, and **** indicates p≤0.0001 for each pairwise comparison.). MH *MAT***a** (XL280**a**) and HH *MAT*α (XL190α) strains, LH *MAT***a** (XL187**a**) and MH*MAT*α (XL280α) strains, NH *MAT***a** (JEC20**a**) and LH *MAT*α (JEC21α) strains, and an enhanced hyphal *MAT*α (JEC21α *gpa3*Δ::*NEO*) strain were grown on V8 agar medium for 36 hours. The expression levels of JEC20**a** or JEC21α were set to 1, and the remaining values of the same mating type strains were normalized to this. The error bars represent the standard deviation of the mean for three biological replicates.

**Figure S4. Exogenous pheromone has modest impact on cell-cell fusion**.

Cell fusion frequencies between *MAT***a** and *MAT*α cells (MH and HH, LH and MH, NH and LH, and NH and EH) co-incubated on (A) V8 and (B) YPD media for four days both in the absence and in the presence of 500 nM α pheromone peptide.

**Figure S5. Enhanced unisexual reproduction results in a fitness cost during vegetative growth**.

Hyphal growth on MS medium for two weeks for *C. neoformans* strains KN99**a**, KN99**a** *gpa3*Δ::*NEO* #1, and KN99**a** *gpa3*Δ::*NEO* #2, and NH, LH, and EH *C. deneoformans* strains JEC20**a**, JEC21α, JEC20**a** *gpa3*Δ::*ADE2 ade2*, and JEC21α *gpa3*Δ::*NEO*. The scale bar represents 500 *µ*m. Growth curves were generated using an automated Tecan Sunrise absorbance reader bi-hourly for 72 hours.

**Figure S6. Experimental strategy for the fitness competition assay**.

Equal number of cells of α LH and α LH strains, and of α LH and α EH strains were mixed and spot-inoculated on both YPD and V8 media both in the absence and in the presence of equal number of **a** MH cells. Cells were scraped off agar medium after 10 days of incubation, and both yeast cells and hyphae were collected.

**Table S1. Bisexual cell fusion frequencies for the mating competition experiment**.

**Table S2. p-Values of one-way ANOVA analyses and Welch’s t-test for each pairwise comparison for the foraging for mating assay during mating confrontation**.

**Table S3. p-Values of one-way ANOVA analyses and Welch’s t-test for each pairwise comparison for the foraging for mating assay during mating among mini-colonies**.

**Table S4. Strains and plasmids used in this study**.

**Table S5. Primers used in this study**.

**Table S6. Mating competition experimental design**.

