## Supplemental figures and tables for "Unisexual reproduction promotes competition for mating partners in the global human fungal pathogen *Cryptococcus deneoformans*"

#### Figure S1

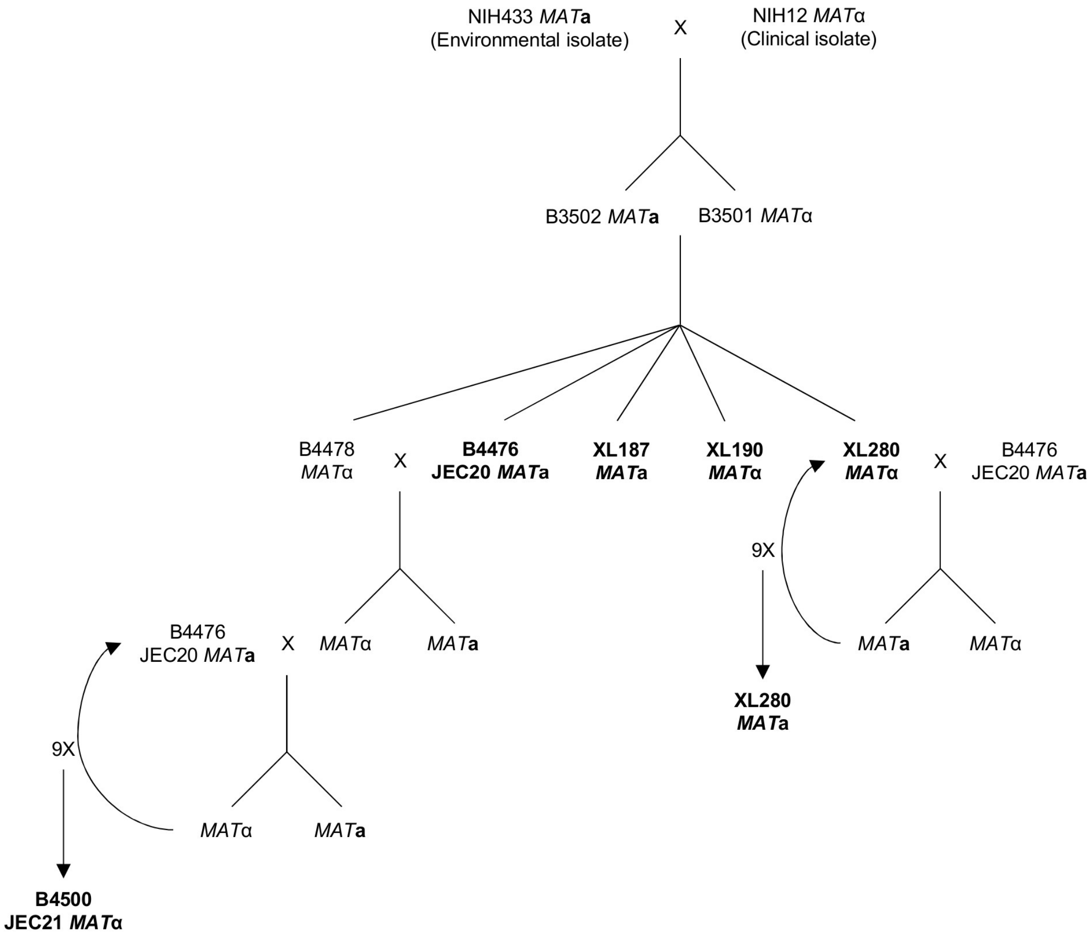
NIH433 MATa
(Environmental isolate)
X
NIH12 MATα
(Clinical isolate)
B3502 MATa
B3501 MATα
B4478
MATα
X
B4476
JEC20 MATa
XL187
MATa
XL190
MATα
XL280
MATα
X
B4476
JEC20 MATa
9X
MATa
MATα
XL280
MATa
B4476
JEC20 MATa
X
MATα
MATa
9X
MATα
MATa
B4500
JEC21 MATα

**Figure S2**

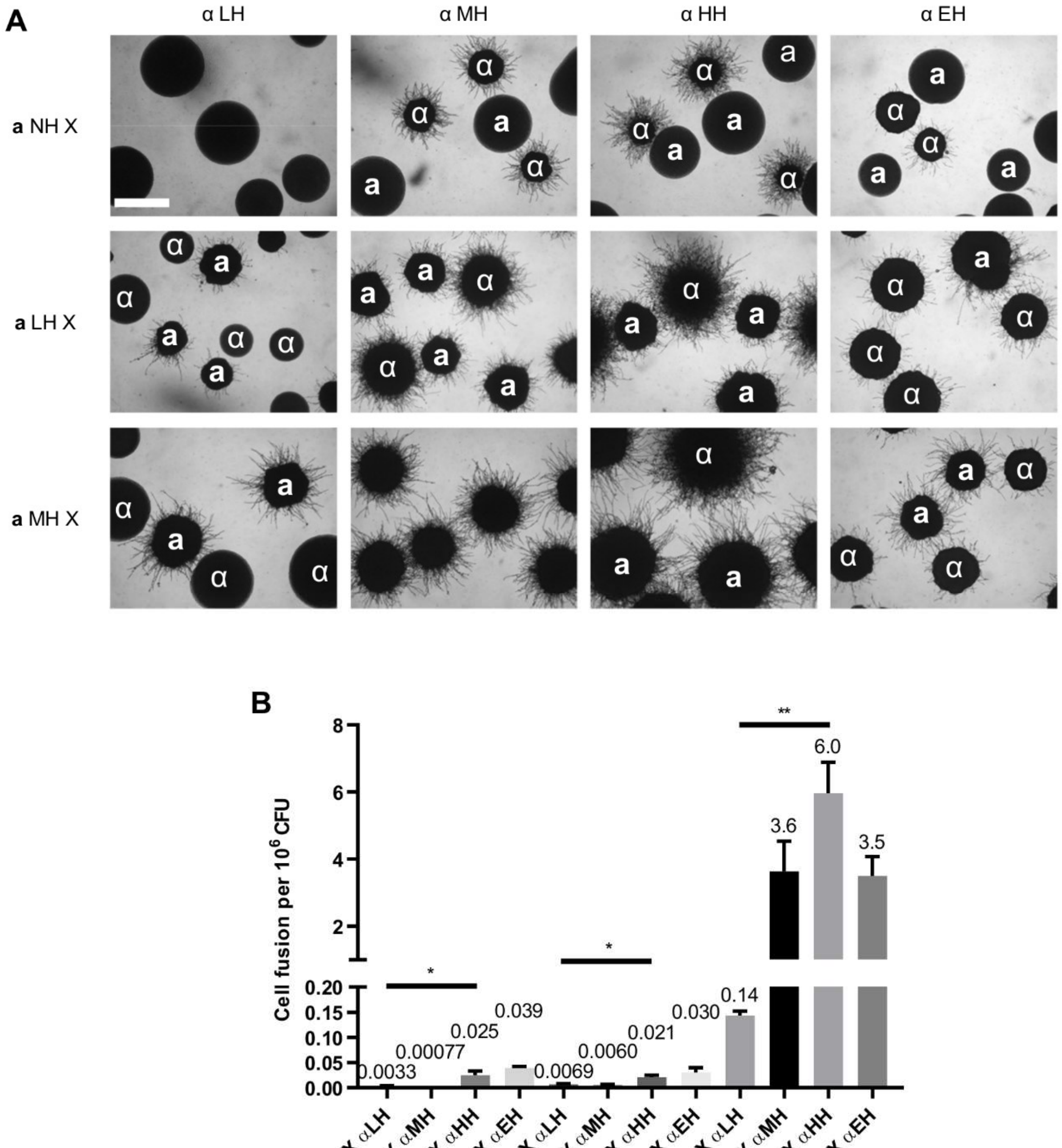
A
α LH
α MH
α HH
α EH
**a** NH X
**a** LH X
**a** MH X
B
Cell fusion per $10^6$ CFU
0.0033
0.00077
0.025
0.039
0.0069
0.0060
0.021
0.030
0.14
3.6
6.0
3.5
*
*
**
a NH X αLH
a NH X αMH
a NH X αHH
a NH X αEH
aLH X αLH
aLH X αMH
aLH X αHH
aLH X αEH
a MH X αLH
a MH X αMH
a MH X αHH
a MH X αEH

### Figure S3

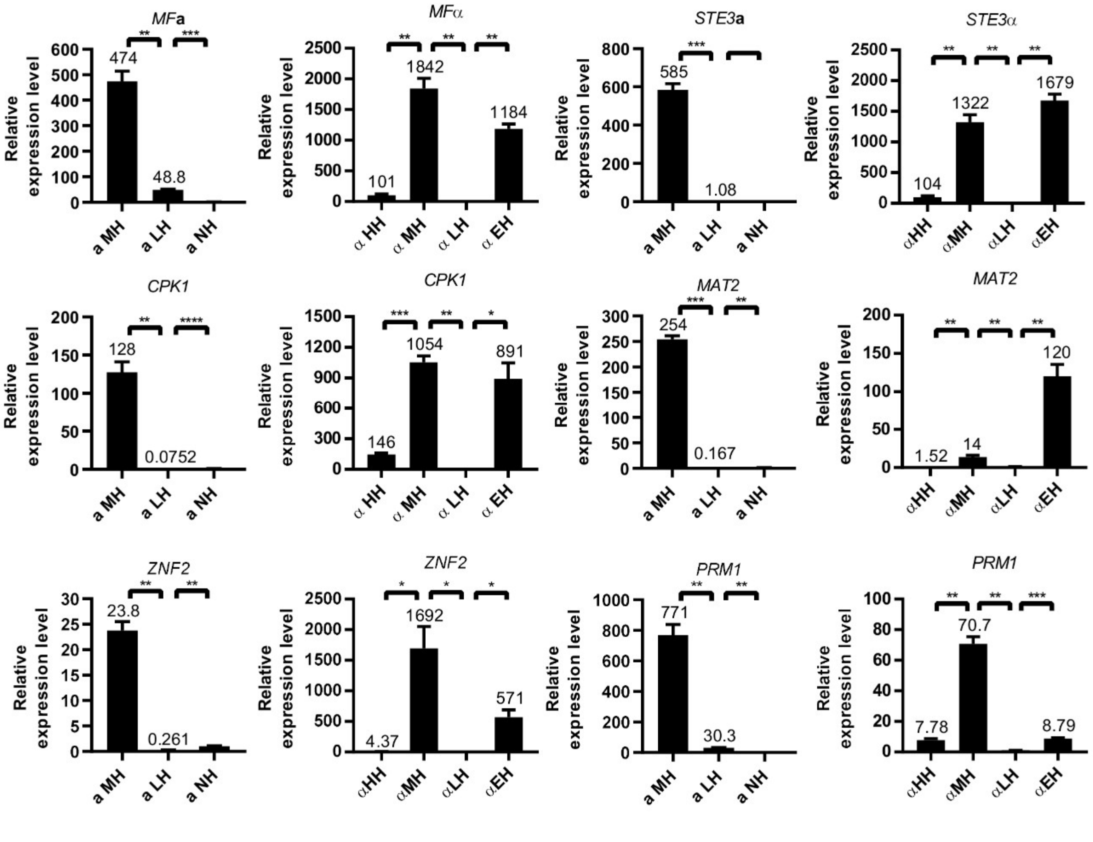

MFa
Relative expression level
474
48.8
a MH
a LH
a NH
MFα
1842
1184
101
α HH
α MH
α LH
α EH
STE3a
585
1.08
STE3α
1322
1679
104
αHH
αMH
αLH
αEH
CPK1
128
0.0752
CPK1
1054
891
146
MAT2
254
0.167
MAT2
120
1.52
14
ZNF2
23.8
0.261
ZNF2
1692
571
4.37
PRM1
771
30.3
PRM1
70.7
7.78
8.79

#### Figure S4

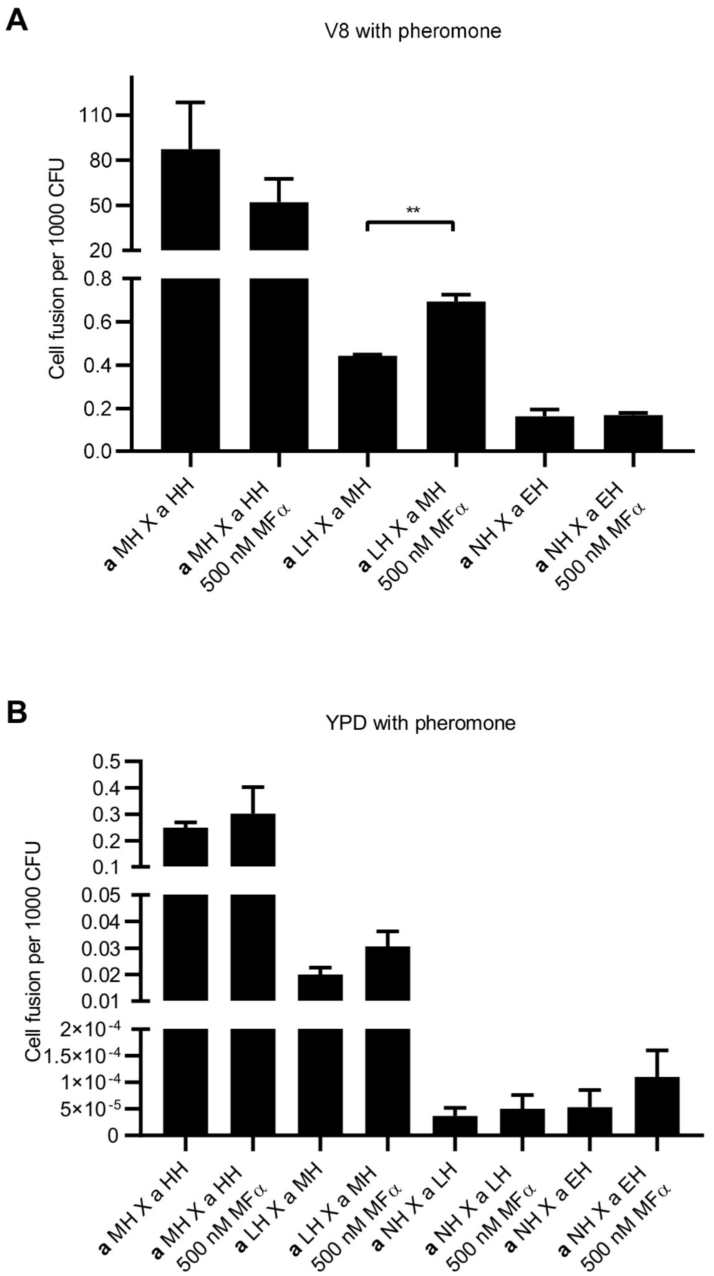
A
V8 with pheromone
Cell fusion per 1000 CFU
110
80
50
20
0.8
0.6
0.4
0.2
0.0
**
a MH X a HH
a MH X a HH 500 nM MFα
a LH X a MH
a LH X a MH 500 nM MFα
a NH X a EH
a NH X a EH 500 nM MFα
B
YPD with pheromone
Cell fusion per 1000 CFU
0.5
0.4
0.3
0.2
0.1
0.05
0.04
0.03
0.02
0.01
2×10-4
1.5×10-4
1×10-4
5×10-5
0
a MH X a HH
a MH X a HH 500 nM MFα
a LH X a MH
a LH X a MH 500 nM MFα
a NH X a LH
a NH X a LH 500 nM MFα
a NH X a EH
a NH X a EH 500 nM MFα

**Figure S5**

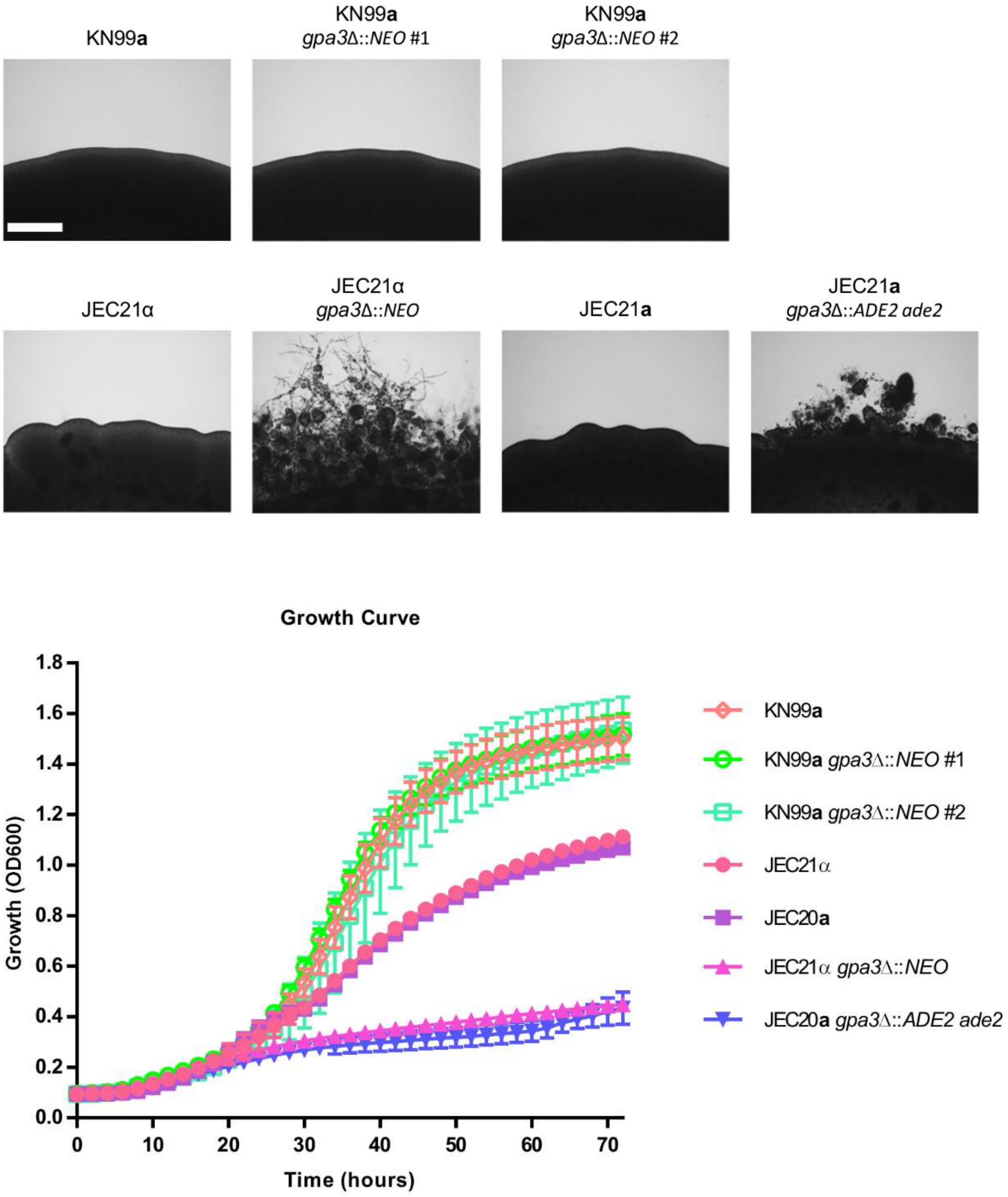

KN99**a**
KN99**a** *gpa3*Δ::*NEO* #1
KN99**a** *gpa3*Δ::*NEO* #2
JEC21α
JEC21α *gpa3*Δ::*NEO*
JEC21**a**
JEC21**a** *gpa3*Δ::*ADE2 ade2*
Growth Curve
Growth (OD600)
Time (hours)
0.0
0.2
0.4
0.6
0.8
1.0
1.2
1.4
1.6
1.8
0
10
20
30
40
50
60
70
KN99**a**
KN99**a** *gpa3*Δ::*NEO* #1
KN99**a** *gpa3*Δ::*NEO* #2
JEC21α
JEC20**a**
JEC21α *gpa3*Δ::*NEO*
JEC20**a** *gpa3*Δ::*ADE2 ade2*

**Figure S6**

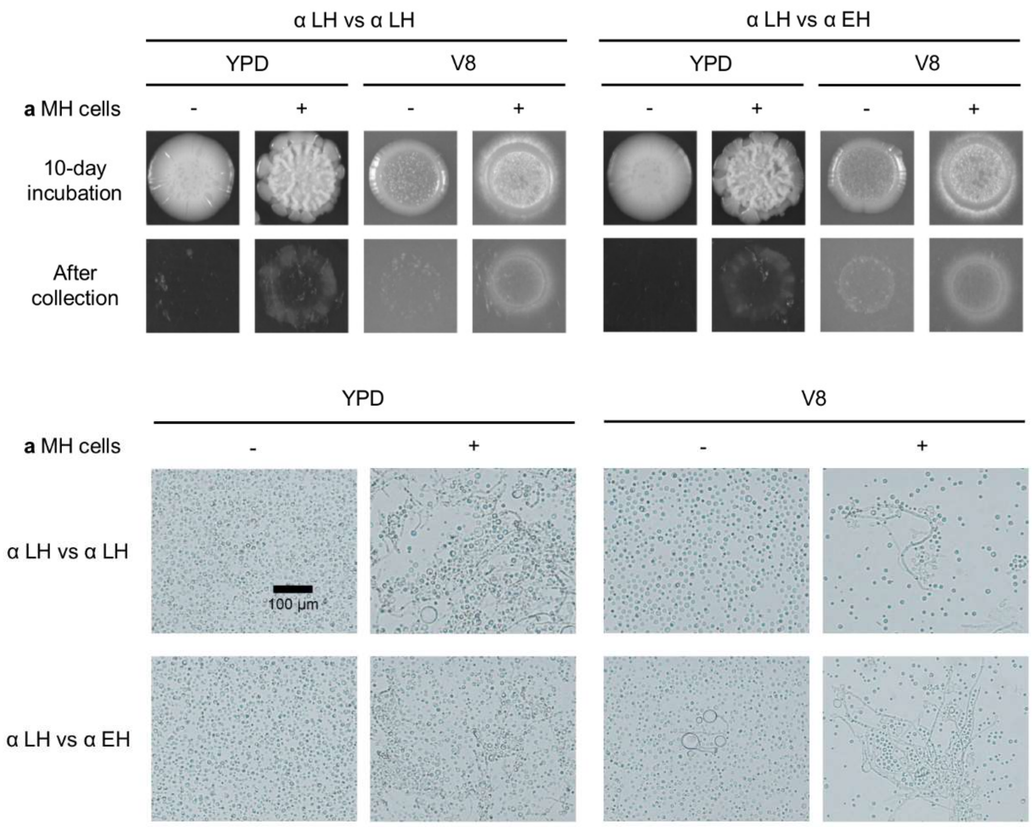
α LH vs α LH
α LH vs α EH
YPD
V8
YPD
V8
a MH cells
-
+
-
+
-
+
-
+
10-day incubation
After collection
YPD
V8
a MH cells
-
+
-
+
α LH vs α LH
100 µm
α LH vs α EH

**Table S1. Bisexual cell fusion frequencies for the mating competition experiment.**

| | Competition pair | Fusion pair | Cell fusion per 1000 CFU | Fold difference | P value |
|---|---|---|---|---|---|
| Competing for **a** MH *HYG* | α-1 HH Vs α-2 MH | α HH X **a** MH | 52 | 8.1 | 0.0005 *** |
| | | α MH X **a** MH | 6.4 | | |
| | α-1 HH Vs α-2 LH | α HH X **a** MH | 10 | 24 | 0.0023 ** |
| | | α LH X **a** MH | 0.41 | | |
| | α-1 HH Vs α-2 LH | α MH X **a** MH | 3.4 | 5.8 | 0.0159 * |
| | | α LH X **a** MH | 0.59 | | |
| Competing for **a** LH *HYG* | α-1 HH Vs α-2 MH | α HH X **a** LH | 1.3 | 6.5 | 0.0003 *** |
| | | α MH X **a** LH | 0.2 | | |
| | α-1 HH Vs α-2 LH | α HH X **a** LH | 0.29 | 14.5 | 0.0228 * |
| | | α LH X **a** LH | 0.02 | | |
| | α-1 HH Vs α-2 LH | α MH X **a** LH | 0.091 | 4.6 | 0.0006 *** |
| | | α LH X **a** LH | 0.02 | | |
| Competing for **a** NH *HYG* | α-1 HH Vs α-2 MH | α HH X **a** NH | 0.049 | 5.3 | 0.0468 * |
| | | α MH X **a** NH | 0.0092 | | |
| | α-1 HH Vs α-2 LH | α HH X **a** NH | 0.064 | 8.9 | 0.0170 * |
| | | α LH X **a** NH | 0.0072 | | |
| | α-1 HH Vs α-2 LH | α MH X **a** NH | 0.01 | 1.7 | 0.0126 * |
| | | α LH X **a** NH | 0.0059 | | |

**Table S2. p-Values of one-way ANOVA analyses and Welch's t-test for each pairwise comparison for the foraging for mating assay during mating confrontation.**

| One way ANOVA group analyses (* 0.01<p≤0.05, ** 0.001<p≤0.01, *** 0.0001<p≤0.001, ****p≤0.0001) | | | | | | | | | | | |
|---|---|---|---|---|---|---|---|---|---|---|---|
| **a** NH X α LH | ** 0.0018 | **a** LH X α LH | * 0.0173 | **a** MH X α LH | **** <0.0001 | **a** NH X α LH | 0.0896 | **a** NH X α MH | **** <0.0001 | **a** NH X α HH | **** <0.0001 |
| **a** NH X α MH | | **a** LH X α MH | | **a** MH X α MH | | **a** LH X α LH | | **a** LH X α MH | | **a** LH X α HH | |
| **a** NH X α HH | | **a** LH X α HH | | **a** MH X α HH | | **a** MH X α LH | | **a** MH X α MH | | **a** MH X α HH | |

| One way ANOVA group analyses (continued) | |
|---|---|
| **a** NH X α EH | **** <0.0001 |
| **a** LH X α EH | |
| **a** MH X α EH | |

| Pairwise Welich's t-test analyses (* 0.01<p≤0.05, ** 0.001<p≤0.01, *** 0.0001<p≤0.001, ****p≤0.0001) | | | | | | | | | | | | |
|---|---|---|---|---|---|---|---|---|---|---|---|---|
| **a** NH X α LH | - | | | | | | | | | | | |
| **a** NH X α MH | * 0.0429 | - | | | | | | | | | | |
| **a** NH X α HH | * 0.0181 | * 0.0382 | - | | | | | | | | | |
| **a** NH X α EH | 0.0591 | - | - | - | | | | | | | | |
| **a** LH X α LH | >0.9999 | - | - | - | - | | | | | | | |
| **a** LH X α MH | - | * 0.0414 | - | - | ** 0.0076 | - | | | | | | |
| **a** LH X α HH | - | - | 0.2937 | | 0.0507 | 0.1324 | - | | | | | |
| **a** LH X α EH | - | - | - | 0.0591 | >0.9999 | - | - | - | | | | |
| **a** MH X α LH | 0.1594 | - | - | - | 0.1594 | - | - | - | - | | | |
| **a** MH X α MH | - | ** 0.0078 | - | - | - | ** 0.0097 | - | - | ** 0.0077 | - | | |
| **a** MH X α HH | - | - | ** 0.0034 | - | - | - | ** 0.0032 | - | ** 0.0029 | * 0.0119 | - | |
| **a** MH X α EH | - | - | - | ** 0.0011 | - | - | - | ** 0.0011 | *** 0.0010 | - | - | - |
| | **a** NH X α LH | **a** NH X α MH | **a** NH X α HH | **a** NH X α EH | **a** LH X α LH | **a** LH X α MH | **a** LH X α HH | **a** LH X α EH | **a** MH X α LH | **a** MH X α MH | **a** MH X α HH | **a** MH X α EH |

**Table S3. p-Values of one-way ANOVA analyses and Welch's t-test for each pairwise comparison for the foraging for mating assay during mating among mini-colonies.**

| One way ANOVA group analyses (* 0.01<p≤0.05, ** 0.001<p≤0.01, *** 0.0001<p≤0.001, ****p≤0.0001) | | | | | | | | | | | | | |
|---|---|---|---|---|---|---|---|---|---|---|---|---|---|
| **a** NH X α LH | * 0.0269 | **a** LH X α LH | * 0.0138 | **a** MH X α LH | ** <0.0043 | **a** NH X α LH | **** <0.0001 | **a** NH X α MH | ** 0.0038 | **a** NH X α HH | *** 0.0003 | **a** NH X α EH | *** 0.0005 |
| **a** NH X α MH | | **a** LH X α MH | | **a** MH X α MH | | **a** LH X α LH | | **a** LH X α MH | | **a** LH X α HH | | **a** LH X α EH | |
| **a** NH X α HH | | **a** LH X α HH | | **a** MH X α HH | | **a** MH X α LH | | **a** MH X α MH | | **a** MH X α HH | | **a** MH X α EH | |

| Pairwise Welich's t-test analyses (* 0.01<p≤0.05, ** 0.001<p≤0.01, *** 0.0001<p≤0.001, ****p≤0.0001) | **a** NH X α LH | **a** NH X α MH | **a** NH X α HH | **a** NH X α EH | **a** LH X α LH | **a** LH X α MH | **a** LH X α HH | **a** LH X α EH | **a MH** X α LH | **a** MH X α MH | **a** MH X α HH | **a** MH X α EH |
|---|---|---|---|---|---|---|---|---|---|---|---|---|
| **a** NH X α LH | | | | | | | | | | | | |
| **a** NH X α MH | * 0.0341 | | | | | | | | | | | |
| **a** NH X α HH | 0.1297 | 0.1081 | | | | | | | | | | |
| **a** NH X α EH | ** 0.0066 | | | | | | | | | | | |
| **a** LH X α LH | * 0.0408 | | | | | | | | | | | |
| **a** LH X α MH | | * 0.0401 | | | 0.5678 | | | | | | | |
| **a** LH X α HH | | | 0.6941 | | 0.0822 | 0.071 | | | | | | |
| **a** LH X α EH | | | | 0.4426 | 0.128 | | | | | | | |
| **a** MH X α LH | ** 0.0038 | | | | ** 0.0038 | | | | | | | |
| **a** MH X α MH | | 0.0565 | | | | 0.0567 | | | 0.0608 | | | |
| **a** MH X α HH | | | * 0.0232 | | | | * 0.0231 | | * 0.0241 | 0.1446 | | |
| **a** MH X α EH | | | | * 0.0267 | | | | * 0.0266 | * 0.0283 | | | |

**Table S4. Strains and plasmids used in this study.**

| | Strain name | Genotype | Background | Sources |
|---|---|---|---|---|
| | XL190α | | | [1] |
| | XL280α | | | [1] |
| | JEC21α | | | [2] |
| | XL280**a** | | | [3] |
| | XL187**a** | | | [4] |
| | JEC20**a** | | | [2] |
| High Hyphal (HH) | XL557 | *NAT* | XL190α | Xiaorong Lin unpublished |
| | CF914 | *NEO* | XL190α | This study |
| Intermediate Hyphal (MH) | CF978 | *HYG* | XL280**a** | This study |
| | CF750 | *NAT* | XL280α | [5] |
| | CF752 | *NEO* | XL280α | [5] |
| | CF974 | *HYG* | XL280α | This study |
| Low Hyphal (LH) | CF931 | *HYG* | XL187**a** | This study |
| | CF969 | *NAT* | JEC21α | This study |
| | CF759 | *NEO* | JEC21α | [5] |
| No Hyphal (NH) | CF926 | *HYG* | JEC20**a** | This study |
| Enhanced Hyphal (EH) | CF1314 | *gpa3Δ::NEO* | JEC21α | This study |
| | CF963 | *crg1Δ::NAT* | XL280α | This study |
| | CF1027 | *gpa3Δ::NEO* | XL280**a** | This study |
| | CF1048 | *crg1Δ::NAT gpa3Δ::NEO* | XL280α | This study |
| | YPH86 | *gpa3Δ::ADE2 ade2* | JEC20**a** | [6] |
| *C. neoformans* strains | KN99**a** | | | [7] |
| | YSB136 | *gpa3Δ::NEO* | KN99**a** | [6] |
| | YSB137 | *gpa3Δ::NEO* | KN99**a** | [6] |
| | **Plasmid** | **Genotype** | **Background** | **Sources** |
| | pAI3 | *NAT AMP* | | [8] |
| | pJAF1 | *NEO AMP* | | [9] |
| | pJAF15 | *HYG AMP* | | [9] |

1. Lin, X., Hull, C.M., and Heitman, J. (2005). Sexual reproduction between partners of the same mating type in *Cryptococcus neoformans*. Nature *434*, 1017-1021.
2. Kwon-Chung, K.J., Edman, J.C., and Wickes, B.L. (1992). Genetic association of mating types and virulence in *Cryptococcus neoformans*. Infect. Immun. *60*, 602-605.
3. Zhai, B., Zhu, P., Foyle, D., Upadhyay, S., Idnurm, A., and Lin, X. (2013). Congenic strains of the filamentous form of *Cryptococcus neoformans* for studies of fungal morphogenesis and virulence. Infect. Immun. *81*, 2626-2637.
4. Lin, X., Huang, J.C., Mitchell, T.G., and Heitman, J. (2006). Virulence attributes and hyphal growth of *C. neoformans* are quantitative traits and the *MAT*α allele enhances filamentation. PLoS Genet. *2*, e187.
5. Fu, C., and Heitman, J. (2017). *PRM1* and *KAR5* function in cell-cell fusion and karyogamy to drive distinct bisexual and unisexual cycles in the *Cryptococcus* pathogenic species complex. PLoS Genet. *13*, e1007113.
6. Hsueh, Y.P., Xue, C., and Heitman, J. (2007). G protein signaling governing cell fate decisions involves opposing Gα subunits in *Cryptococcus neoformans*. Mol. Biol. Cell *18*, 3237-3249.
7. Nielsen, K., Cox, G.M., Wang, P., Toffaletti, D.L., Perfect, J.R., and Heitman, J. (2003). Sexual cycle of *Cryptococcus neoformans* var. *grubii* and virulence of congenic **a** and α isolates. Infect. Immun. *71*, 4831-4841.
8. Idnurm, A., Reedy, J.L., Nussbaum, J.C., and Heitman, J. (2004). *Cryptococcus neoformans* virulence gene discovery through insertional mutagenesis. Eukaryot. Cell *3*, 420-429.
9. Fraser, J.A., Subaran, R.L., Nichols, C.B., and Heitman, J. (2003). Recapitulation of the sexual cycle of the primary fungal pathogen *Cryptococcus neoformans* var. *gattii*: implications for an outbreak on Vancouver Island, Canada. Eukaryot. Cell *2*, 1036-1045.

**Table S5. Primers used in this study.**

| Primer name | Sequence (5' to 3') | Description |
|---|---|---|
| M13F | GTAAAACGACGGCCAGT | *NAT*/*NEO* cassette |
| M13R | CAGGAAACAGCTATGAC | *NAT*/*NEO* cassette |
| JOHE41446 | GCCGTGCAAGGGTGTAGG | *SXI1*α F |
| JOHE41447 | GGGCCATTGGAGGAAGCTG | *SXI1*α R |
| JOHE41444 | CGGACGAGCTCTCAAATTGG | *SXI2***a** F |
| JOHE41445 | TTTGCTCGCTCTCCTTCCAC | *SXI2***a** R |
| JOHE43073 | CATTGAAACTCCCTGCTTGG | *GPA3* 5'UTR F |
| JOHE43075 | ACTGGCCGTCGTTTTACCGTCTGAAAGT TGGTCGTTG | *GPA3* 5'UTR R |
| JOHE43077 | GTCATAGCTGTTTCCTGTCTCTCTGTGG CTCGATTT | *GPA3* 3'UTR F |
| JOHE43078 | GGAACTCGCCCTCAATCTC | *GPA3 3*'UTR R |
| JOHE43072 | GCAAGAAGAGGTGAGCAGTC | *GPA3* Junction F |
| JOHE43079 | ACGTTCGTAAAGGGGTTGG | *GPA3* Junction R |
| JOHE43074 | CCGAGCATCAGACGAACAC | *GPA3* F |
| JOHE43076 | AATAGCGAGACGCACATCC | *GPA3* R |
| JOHE43065 | CGGGTTGCTTTATCTCGTTC | *CRG1* 5'UTR F |
| JOHE43067 | ACTGGCCGTCGTTTTACTTATCCCAGGC AGCGTTCT | *CRG1* 5'UTR R |
| JOHE43069 | GTCATAGCTGTTTCCTGTCGCTTCTTTC CCCGATCTAC | *CRG1* 3'UTR F |
| JOHE43070 | AGAGGCTTCGGCAAGATCAT | *CRG1 3*'UTR R |
| JOHE43064 | TTTCCCTTCTGTCCCCATC | *CRG1* Junction F |
| JOHE43071 | TCGAGATGCTGGTAGGCACA | *CRG1* Junction R |
| JOHE43066 | TCTTCTTCTCTCTCGCCTCCT | *CRG1* F |
| JOHE43068 | GAATGTCGTAGTGGTCGTGGT | *CRG1* R |
| JOHE44120 | GTCTCCACTGATTTCATTGGCTCTAC | *GPD1* RTPCR F |
| JOHE44121 | GTAACCATACTCATTGTCATACCAGCTG | *GPD1* RTPCR R |
| JOHE43005 | ATCTTCACCACCTTCACTTCT | *MF*α RTPCR F |
| JOHE43006 | CTAGGCGATGACACAAAGG | *MF*α RTPCR R |
| JOHE45716 | GGACGCCTTCACTGCTATCT | *MF***a** RTPCR F |
| JOHE45717 | GCTACCGTAAGCCTCTTCGTT | *MF***a** RTPCR R |
| JOHE44039 | CTCCTTGTCCTTTTACCTCTGC | *STE3*α RTPCR F |
| JOHE44040 | CTTGTGGCTGAAATCCCAAC | *STE3*α RTPCR R |
| JOHE45718 | GACGGTATCACTGGTTGTCT | *STE3***a** RTPCR F |
| JOHE45719 | CGTACTATCCTCGCTTCATC | *STE3***a** RTPCR R |
| JOHE44033 | CAGGTTCAACGTCGGCAACAACTA | *CPK1* RTPCR F |

| | | |
|---|---|---|
| JOHE44034 | TCAAGTCGCGATGGATGATTTCAGCAG | *CPK1* RTPCR R |
| JOHE44027 | AGGAAGCTCGAGCATCGAAAGCTGTAT | *MAT2* RTPCR F |
| JOHE44028 | TGAGCTGCAGTAGCTCGTAAATCTGAC | *MAT2* RTPCR R |
| JOHE44029 | ACCCTTTCCAACCCCTTGTCAACG | *ZNF2* RTPCR F |
| JOHE44030 | AAGGACGTTTCCCAGTGTGAATACGCC | *ZNF2* RTPCR R |
| JOHE42831 | TGCCTCTTCTTCCGTATCGT | *PRM1* RTPCR F |
| JOHE42832 | CCCCAAAGGGATCTTTTCTC | *PRM1* RTPCR R |

**Table S6. Mating competition experimental design.**

| | Unisex | #1 α | #2 α | |
|---|---|---|---|---|
| #1 | HH X HH | XL190α *NAT* | XL190α *NEO* | |
| #2 | MH X MH | XL280α *NAT* | XL280α *NEO* | |
| #3 | LH X LH | JEC21α *NAT* | JEC21α *NEO* | |
| | Bisex | #1 α | #2 **a** | |
| #4 | HH X MH | XL190α *NAT* | XL280**a** *HYG* | |
| #5 | MH X LH | XL280α *NAT* | XL187**a** *HYG* | |
| #6 | LH X NH | JEC21α *NAT* | JEC20**a** *HYG* | |
| | Unisex competition | #1 α | #2 α | #3 α |
| #7 | MH/HH/LH | XL280α *HYG* | XL190α *NAT* | JEC21α *NEO* |
| | Bisex competition | #1 α | #2 α | #3 **a** |
| #8 | MH/HH/MH | XL280α *NAT* | XL190α *NEO* | XL280**a** *HYG* |
| #9 | MH/LH/MH | XL280α *NAT* | JEC21α *NEO* | XL280**a** *HYG* |
| #10 | HH/LH/MH | XL190α *NAT* | JEC21α *NEO* | XL280**a** *HYG* |
| #11 | MH/HH/LH | XL280α *NAT* | XL190α *NEO* | XL187**a** *HYG* |
| #12 | MH/LH/LH | XL280α *NAT* | JEC21α *NEO* | XL187**a** *HYG* |
| #13 | HH/LH/LH | XL190α *NAT* | JEC21α *NEO* | XL187**a** *HYG* |
| #14 | MH/HH/NH | XL280α *NAT* | XL190α *NEO* | JEC20**a** *HYG* |
| #15 | MH/LH/NH | XL280α *NAT* | JEC21α *NEO* | JEC20**a** *HYG* |
| #16 | HH/LH/NH | XL190α *NAT* | JEC21α *NEO* | JEC20**a** *HYG* |
| | Unisex | #1 α | #2 α | |
| #17 | LH/LH | JEC21α *NEO* | JEC21α *NAT* | |
| #18 | IH/LH | JEC21α *gpa3::NEO* | JEC21α *NAT* | |
| | Unisex competition | #1 α | #2 α | #3 α |
| #19 | LH/LH/MH | JEC21α *NEO* | JEC21α *NAT* | XL280α *HYG* |
| #20 | IH/LH/MH | JEC21α *gpa3::NEO* | JEC21α *NAT* | XL280α *HYG* |

| | Bisex | #1 α | #2 α | |
|---|---|---|---|---|
| #21 | LH/NH | JEC21α *NEO* | JEC20**a** *HYG* | |
| #22 | IH/NH | JEC21α *gpa3::NEO* | JEC20**a** *HYG* | |
| | Bisex competition | #1 α | #2 α | #3 **a** |
| #23 | LH/LH/NH | JEC21α *NEO* | JEC21α *NAT* | JEC20**a** *HYG* |
| #24 | IH/LH/NH | JEC21α *gpa3::NEO* | JEC21α *NAT* | JEC20**a** *HYG* |
